# Genetic features and genomic targets of human KRAB-Zinc Finger Proteins

**DOI:** 10.1101/2023.02.27.530095

**Authors:** Jonas de Tribolet-Hardy, Christian W. Thorball, Romain Forey, Evarist Planet, Julien Duc, Bara Khubieh, Sandra Offner, Jacques Fellay, Michael Imbeault, Priscilla Turelli, Didier Trono

## Abstract

Krüppel-associated box (KRAB) domain-containing zinc finger proteins (KZFPs) are one of the largest groups of transcription factors encoded by tetrapods, with 378 members in human alone. KZFP genes are often grouped in clusters reflecting amplification by gene and segment duplication since the gene family first emerged more than 400 million years ago. Previous work has revealed that many KZFPs recognize transposable element (TE)-embedded sequences as genomic targets, and that KZFPs facilitate the co-option of the regulatory potential of TEs for the benefit of the host. Here, we present a comprehensive survey of the genetic features and genomic targets of human KZFPs, notably completing past analyses by adding data on more than a hundred family members. General principles emerge from our study of the TE-KZFP regulatory system, which point to multipronged evolutionary mechanisms underlaid by highly complex and combinatorial modes of action with strong influences on human speciation.

## INTRODUCTION

Krüppel-associated box (KRAB) domain-containing zinc finger proteins (KZFPs) constitute one of the largest groups of transcription factors encoded by tetrapods, with 378 protein-coding representatives in human alone (Table S1). KZFPs are characterized by an N-terminal KRAB domain and a C-terminal array of zinc fingers (ZF) conferring sequence-specific polynucleotide binding potential. KZFP genes are often grouped in clusters reflecting their amplification by gene and segment duplication since the family first emerged more than 400 million years ago in the last common ancestor of lung fish, coelacanth and tetrapods (Huntley et al. 2006; Nowick et al. 2010; Imbeault, Helleboid, and Trono 2017). The KRAB domain of all evolutionarily recent human KZFPS recruits KAP1 (KRAB-associated protein 1), also known as TRIM28 (tripartite motif protein 28), which acts as a scaffold for a heterochromatin-inducing complex repressing transcription over KZFP-bound loci and flanking regions (Urrutia 2003). Older, more conserved KZFPs often harbor variant KRAB domains that display functionally diverse KAP1-devoid protein interactomes (Helleboid et al. 2019). Cumulated work has identified transposable elements (TEs) as major targets of KZFPs, which likely evolved both to control the spread of these genetic invaders and to facilitate the domestication of their regulatory potential (Wolf and Goff 2009; Jacobs et al. 2014; Najafabadi et al. 2015; Schmitges et al. 2016; Imbeault et al. 2017; Bruno et al. 2019). TE-derived sequences make up a readily recognizable 50% of the human genomic DNA, a likely underestimation of their real contribution to our genetic makeup as their signature features get lost over time due to genetic drift. Most human TEs are retrotransposons spreading by a copy-and-paste mechanism, be they LTR (long terminal repeat)-containing endogenous retroviruses (ERVs), long and short interspersed nuclear elements (LINEs and SINEs), or the composite SINE-VNTR-Alu (SVAs). ERVs and LINEs encode the reverse transcriptase and endonuclease activities necessary for their retrotransposition, whereas the non-autonomous SINEs and SVAs elements rely on LINE proteins for spreading. Due respectively to internal recombination and abortive retrotranscription, incomplete ERV and LINE integrants abound, for the former as solo-LTRs and for the latter as 5’-deleted units of various lengths (Kojima 2018).

Uncontrolled TE activity is deleterious to an organism notably because new insertions can disrupt the genome and cause disease (Hancks and Kazazian 2016; Durnaoglu et al. 2021; Kim et al. 2016). Accordingly TEs are tamed by several general mechanisms, whether proteinbased repressors such as the KZFP/KAP1 or the HUSH complexes (Seczynska et al. 2021) or RNA-based mechanisms such as piRNAs (Ozata et al. 2019), which play a prominent role in controlling TEs during the genome reprogramming associated with gametogenesis. However, cumulated evidence indicates that KZFPs do more than simply preventing transposition, be it only because most TE integrants remain targeted by these proteins many millions of years after they have lost all replicative potential due to mutations (Imbeault et al. 2017). This has led to the suggestion that KZFPs act to facilitate the domestication of the regulatory potential of TEs, which had been proposed long ago to be key to the establishment and evolution of gene regulatory networks (Trono 2015; Britten and Davidson 1969). In line with this hypothesis, individual human KZFPs have been found to be implicated in a variety of biological processes, including embryonic genome activation, gastrulation, gametogenesis, imprinting, placentation, brain development, adipogenesis and angiogenesis (Chen et al. 2019; Playfoot et al. 2021; Pontis et al. 2019, 2022; Hayashi and Matsui 2006; Takahashi et al. 2019; Quenneville et al. 2011; Wagner et al. 2000; Zeng et al. 2012; Yang et al. 2017a; Wang et al. 2022; Iouranova et al. 2022; Li et al. 2008). An important step towards defining the roles of all human KZFPs lies in a more complete characterization of their genomic targets. Previous studies have provided a significant advance in this direction, identifying the binding preference of some 240 KZFPs (Helleboid et al. 2019; Imbeault et al. 2017; Najafabadi et al. 2015). Here we complete this effort by unveiling the binding loci of an additional 110 human KZFPs. With these and previously obtained data we now have identified the genomic targets of about 95% human KZFPs, which together with an examination of genetic features of this gene family allows one to delineate some interesting general principles.

## RESULTS

### Genomic distribution and evolutionary features of human KZFP genes

As a starting point, we updated the census of human KZFP genes, using hg19 as data source. We identified 467 pairs of neighbouring sequences corresponding to KRAB and C2H2 polyzinc finger domains, 378 of which were predicted to encode full-length KZFPs. We could also delineate 31 clusters, that is, groups of at least three genes separated by less than 250 kb as previously defined (Huntley et al. 2006) (Figure 1A and Table S1). Eleven of these clusters reside on chromosome 19, collectively hosting 246 KZFPs (219 of which are protein-coding). Using previously estimated evolutionary ages (Imbeault et al. 2017), we further determined that isolated protein-coding KZFP genes tend to be older than their cluster-associated counterparts (p = 3.5e-6, Wilcoxon rank-sum test [WRS]). This fits with the proposal that chromosome 19 is the main region of emergence of new KZFP genes (Lukic et al., 2014) and suggests that escaping the tumultuous environment of this chromosome facilitated the fixation of older family members. However, this is not a strict rule, as the long arm of chromosome 7 hosts both a cluster of some of the most ancient KZFPs (*ZNF282, ZNF777*, and *ZNF783*) near its distal end as previously noted (Liu et al. 2014), and a cluster of primate-specific KZFPs (*AC115220.1, ZNF727, ZNF735, ZNF679, ZNF736, ZNF680, ZNF107, ZNF138, ZNF273 and ZNF117*) near the centromere (Figure 1A and S1).

**Figure 1:**
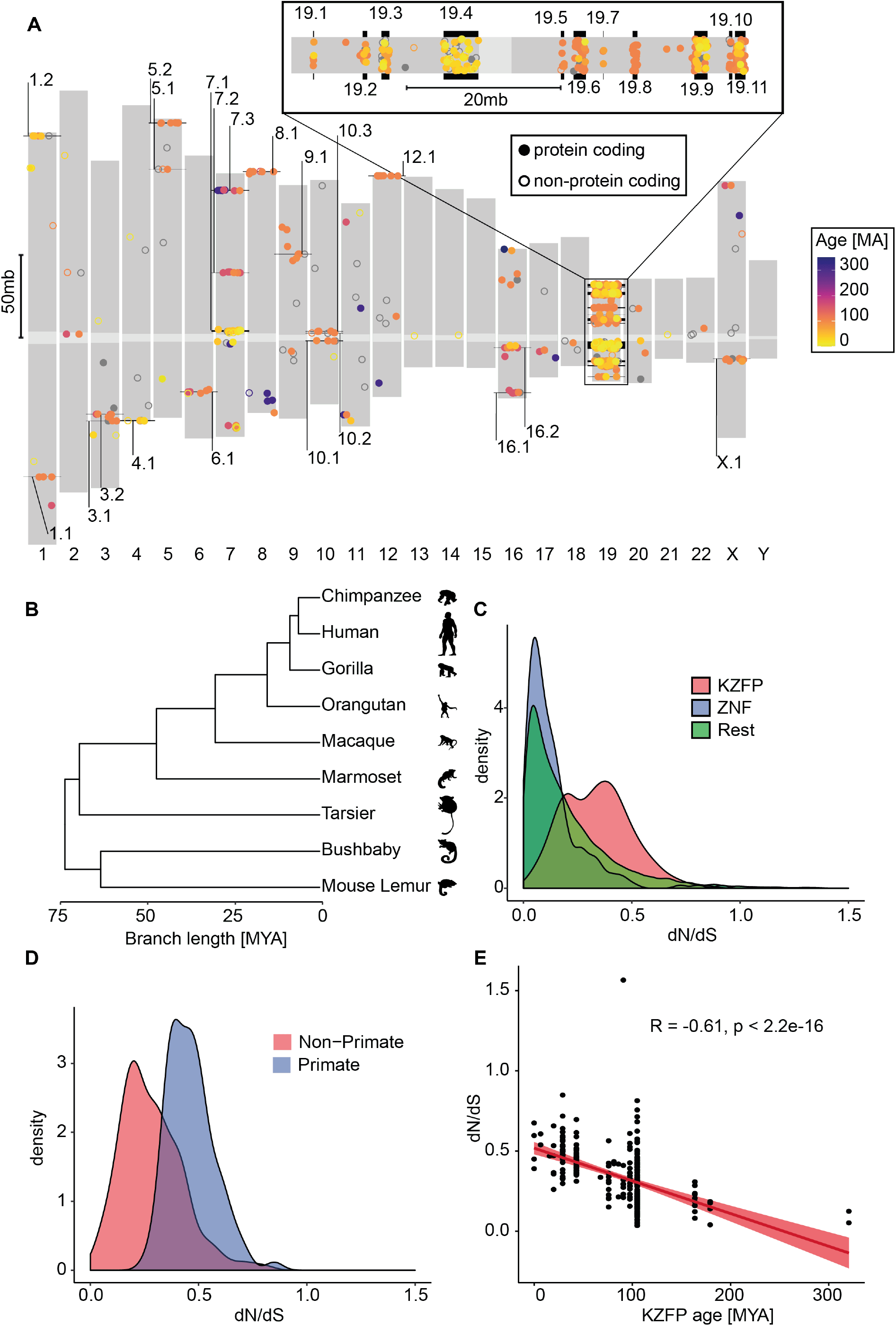
Human KRAB-Zinc Finger Proteins (KZFP) and their evolution in the primate lineage. A) Dots indicate relative chromosomal position of KZFP genes (defined by juxtaposed KRAB- and zinc finger-coding domains), with the colour code indicative of age (grey for unassigned) and numbered clusters pointed to in black. Non-protein coding genes are indicated by a hollow circle. Higher magnification of chromosome 19 is presented on top. Centromeres are indicated in light grey B) Phylogenetic tree of primate species used to calculate natural selection of human genes, with branch length indicating approximate time of divergence in million years (MYA). Silhouettes courtesy of http://phylopic.org/. C) Distribution of PAML dN/dS values of natural selection for KZFPs (red), non-KRAB ZFPs (blue) and all remaining genes in the genome (gray). D) dN/dS distribution of primate-specific KZFPs (blue) and older (red) KZFP genes. E) Spearman correlation of the dN/dS values and estimated age of KZFP genes. The linear regression and 95% confidence interval are shown in red.

To complement this initial analysis, we examined the recent evolution of KZFP genes in the primate lineage as previously described (Takahashi et al. 2019). For this, we determined their gene-wide ratio of non-synonymous (missense) (dN) to synonymous (dS) substitutions (dN/dS), based on genome sequence data from human, chimpanzee, gorilla, orangutan, macaque, marmoset, tarsier, galago (a.k.a. bush baby) and mouse lemur, that is, over ~6 to ~74 million years of divergence (Figure 1B). We found KZFPs to have significantly higher dN/dS values than genes coding for other proteins (p = 1.46e-49, WRS), including KRAB-less ZF proteins (p = 3.04e-46, WRS) (Figure 1C), confirming previous observations (Emerson and Thomas 2009; Najafabadi et al. 2017). We also noted that the distribution of dN/dS values was bimodal amongst KZFPs, with younger, primate-specific genes displaying higher scores than family members having emerged earlier in evolution (p = 2.69e-21, WRS) (Figure 1D) and with the dN/dS ratio of KZFP genes anti-correlated with their estimated age (rho = −0.61) (Figure 1E).

### Conservation-related gradient of human KZFP genes polymorphism

The coding constraint of a gene or fragment thereof reflects the strength of selective pressures imposed on its sequence, hence is linked to the relative functional importance of the corresponding protein or protein domain for a given species. Typically, highly constrained coding regions correspond to loci where mutations are either associated with disease or are completely absent because they cause sterility or embryonic lethality. To calculate the coding constraints imposed on human KZFP genes, we examined genetic variation amongst 138,632 individuals (15,496 genomes and 123,136 exomes) catalogued in gnomAD v.2.0.2 (https://gnomad.broadinstitute.org/). After removing coding sequences with low coverage and dismissing singletons to reduce the impact of false positives resulting from sequencing or alignment errors, we extracted protein-altering variants (missense and predicted loss of function -LoF- by frameshift, gain of stop codon or alteration of essential splice sites) within the canonical transcripts of all remaining KZFPs (n = 361). For the estimation of gene-wide constraint, we normalized the number of variants for the length of the canonical coding sequence and translated the result into a z-score to standardize values (Figure 2A and B). Accordingly, negative deviation from the mean is a sign of increased purifying selection as a consequence of reduced frequency of protein-altering variants. However, we did not correct for a theoretically expected number of mutations as frequently done in this type of analysis because the unstable structure of the ZF array-coding region of KZFP genes renders this parameter unpredictable (Yang et al. 2017b). Gene-wide, LoF and ZF domain-specific scores modestly correlated with previously measured dN/dS ratios and with the age of the KZFPs (Figure 2C). Examining individual domains revealed that this association stemmed mainly from the ZF C2H2- and to a lesser extent fingerprint-coding sequences. Of note, other codons of the ZF-coding regions displayed no significant constraint, confirming that essential positions in ZFs are limited to the structure-conferring cysteine and histidine residues and the target-defining fingerprint residues at positions −1, +3 and +6 of the ZF alpha helix (Najafabadi et al. 2017). Primate-restricted, younger KZFPs were significantly less constrained both in terms of LoF (p = 9.1e-13, WRS) and missense variation (p = 1.7e-6, WRS) than their older counterparts (Figure 2D), with the difference mainly residing in sequences coding the poly-ZF (p = 4.6e-9, WRS) rather than the KRAB domain (p = 0.25, WRS). Within ZFs, the C2H2- and fingerprint-defining positions were again the most influential (p_ZFc2h2_ = 1.1e-14 and p_ZFprint_ = 7.9e-7, WRS), compared to the other non-functional positions of the ZF domains (p_ZF other_ = 0.002, WRS). Correlating with their age, isolated KZFP genes were more constrained at sequences encoding the ZF C2H2 residues (p = 0.001, WRS) and fingerprint-defining positions (p = 0.02, WRS). Furthermore, they displayed lower LoF scores (p = 0.001, WRS) than their cluster-associated counterparts, consistent with their stabilization over longer evolutionary times (Figure 2E). However, coding constraints were also highly heterogeneous within most clusters, indicating that differential selective pressures are rapidly exerted on members of a same gene cluster (Figure S2).

**Figure 2:**
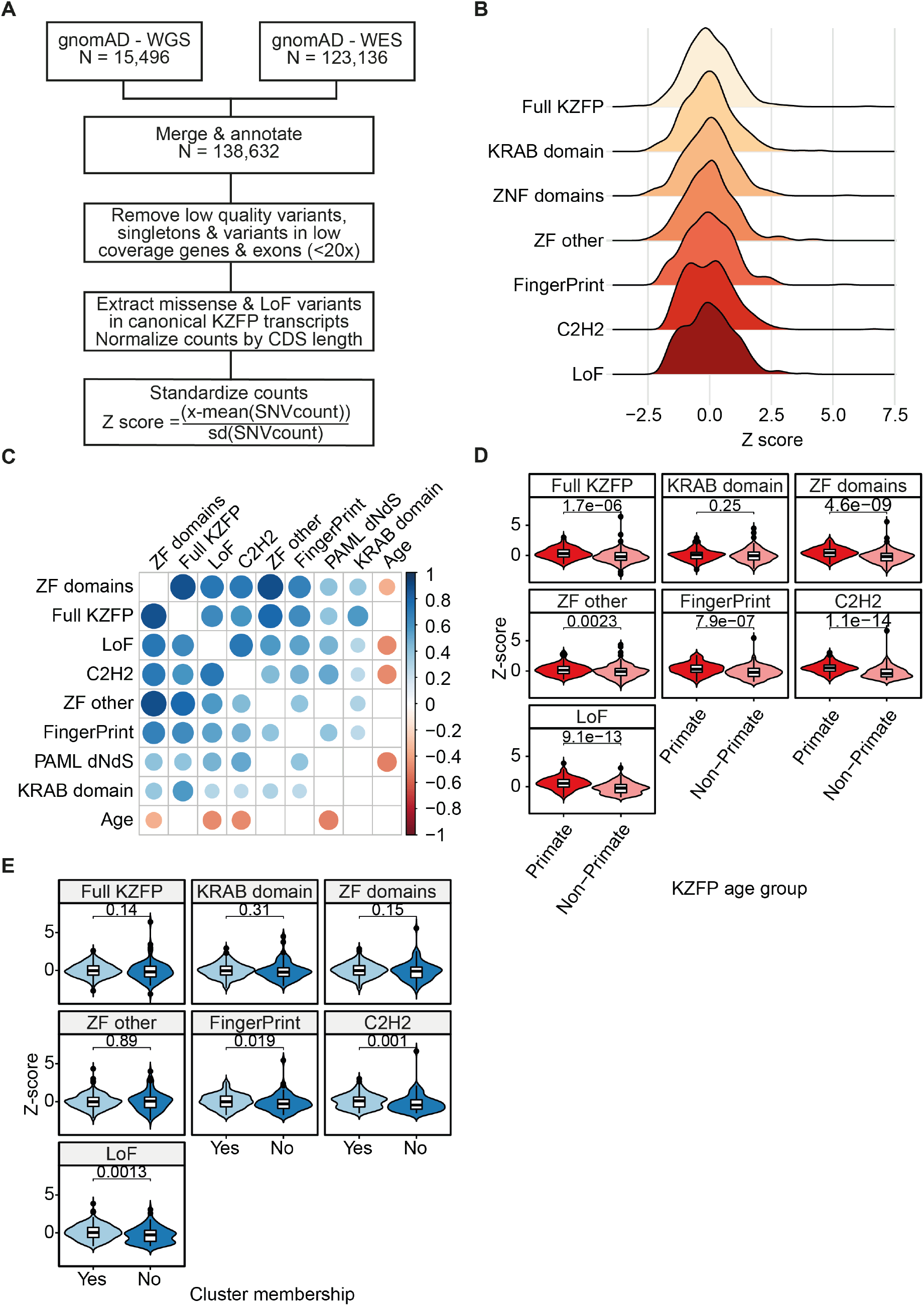
Coding constraints of KZFP genes. A) Schematic of genetic constraint z-score calculation. WGS/WES, whole genome/exome sequencing; LoF, loss of function variant; CDS, coding sequence. B) Distribution of indicated z-scores; a lower score indicates increased constraint compared to the average of all KZFPs. Full KZFP, all variants within the canonical KZFP transcript; KRAB domain, only variants in the KRAB domain; ZF domains, variants within the ZF domains; ZF other, variants in nonfunctional positions within the ZF domains; FingerPrint, variants in the ZF fingerprint positions; C2H2, variants in the cysteine or histidine positions of the ZF domains; LoF, loss of function variants C) Correlation plot showing the Spearman correlations between: the Z-scores defined in Panel B, level of natural selection (PAML dN/dS) and estimated age of KZFPs. The colors and their intensity represent the direction and strength of the correlations, with blue representing a positive- and red a negative correlation. Only significant correlations after Bonferroni correction are shown. D) Primate vs. non-primate KZFP constraint across indicated KZFP domains or residues. E) Relative 552 constraint of indicated regions for KZFPs inside vs. outside clusters. P-values were calculated using the Wilcoxon rank-sum test (WRS).

When looking at the most conserved KZFPs, no LoF variants were detected amongst all examined individuals for *ZFP92, ZNF606, ZNF81, ZNF777, ZNF250*, and *ZNF597*, which all are 105 myo except ZNF777 which is 312 myo. The *ZFP92, ZNF81*, and *ZNF777* genes were also devoid of any missense mutations in their C2H2- or fingerprint-coding positions, while some were detected in *ZNF250, ZNF597*, and *ZNF606* albeit at extremely low allele frequencies. For a majority of other KZFPs (n = 213), some heterozygous but no homozygous LoF variants were observed. Nevertheless, a significant number (n = 148) presented homozygous LoF variants in at least two individuals, suggesting reduced constraint (p < 2.22e-16, WRS) (Figure S2). On average, members of this subgroup had a younger estimated age than the rest of the KZFPs (p = 2.9e-13, WRS).), showing that KZFPs which are more conserved during evolution also have higher constraint in the human population.

### Differential coding constraints of human KZFP paralogs

In order to investigate further the connection between the coding constraints and the evolutionary history of KZFP genes, we examined 33 sets of KZFP paralogs, identified based on similarities between their zinc fingerprints (Imbeault et al. 2017). For 28 of them, both members of a paralog pair were located within the same chromosomal cluster. Significant differences were noted, especially at the C2H2-coding positions, with some pairs of paralogs displaying closely similar coding constraints (e.g. *ZNF75A* and *ZNF75D*) while others were markedly divergent (e.g. *ZNF160* and *ZNF665*) (Figure 3A). The level of divergence was not related to the age of the paralog pairs (p = 0.21, WRS). However, the more constrained paralog within a pair was usually also the most conserved in evolution (Figure 3A). For instance, the ~90-million-year old (myo) *ZNF160* was markedly more constrained than its ~29 myo *ZNF665* paralog, both at C2H2-coding positions and across other features (Figure 3B). A closer examination of *ZNF160* and *ZNF665* zinc fingerprints revealed that some ZFs were completely constrained in both KZFPs, while others were more flexible (Figure 3C). ChIP-seq analyses confirmed that these proteins recognized closely related sequence motifs (Figure 3D) in overlapping sets of genomic targets (Figure 3E), notably some LINE1 integrants (Figure S3A). Furthermore, the two paralogs were noted to have roughly similar expression patterns of across 40 tissues according to the GTEx database (GTEx Consortium et al. 2017) (rho = 0.89) (Figure S3B). This is contrasted with the more global observation that more constrained KZFPs were generally expressed at higher levels and more ubiquitously than their more flexible counterparts (Figure S3C), in line with previous reports (Lek et al. 2016).

**Figure 3:**
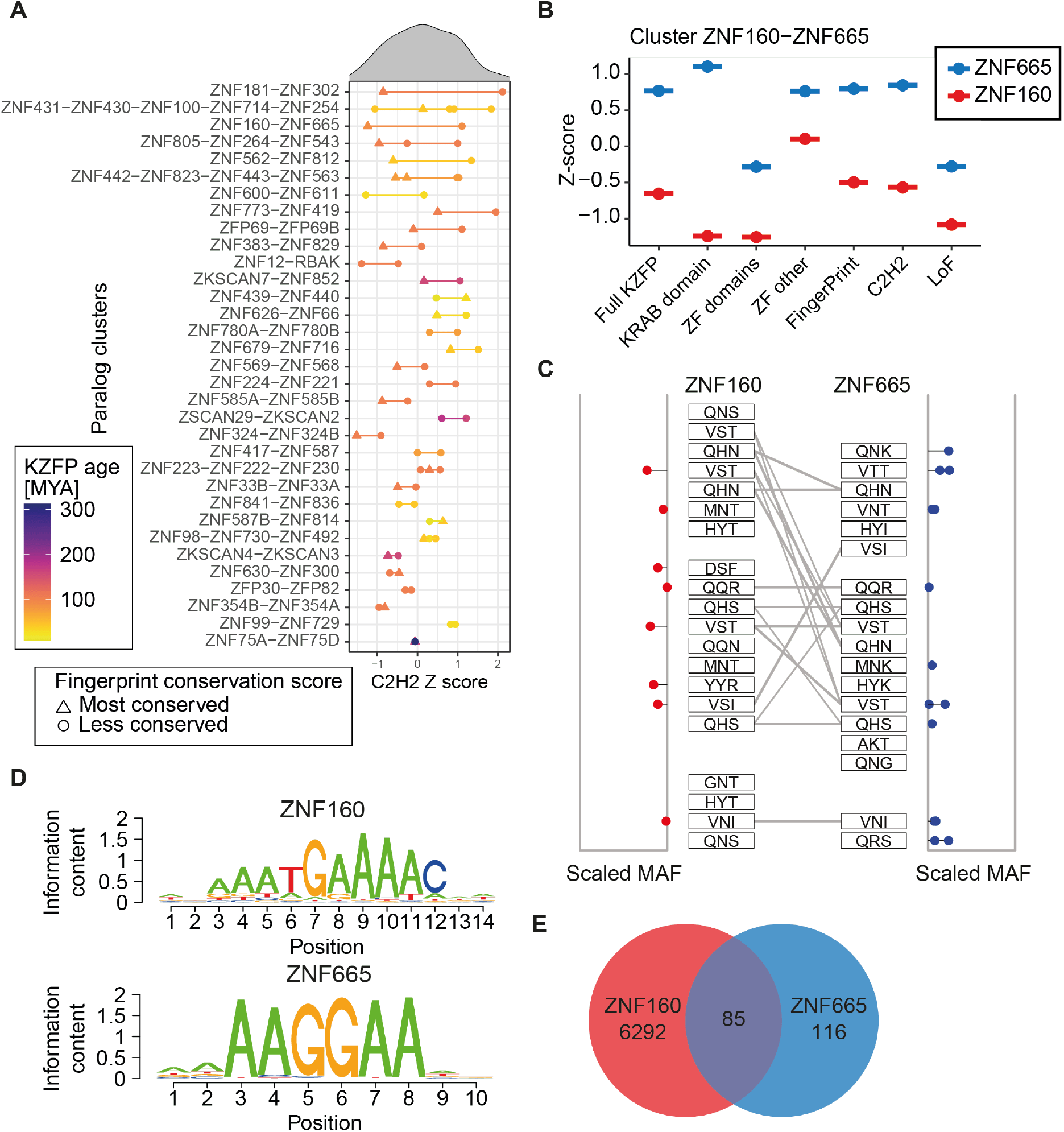
Coding constraints of KZFP paralogs. (A) Distribution of C2H2 constraint z-scores for indicated sets of KZFP paralogs, arranged from top to bottom according to difference within pairs. Each KZFP is colored according to their respective age, with the line separating them colored as the mean age of the pair. The paralog within each pair with the most conserved fingerprint across evolutionary time is marked by a triangle, while less or identically conserved KZFPs are marked by a dot. The order of the y-axis labels corresponds to the order of the colored points on the graph. (B) Differential constraint z-scores for indicated domains of paralogs ZNF160 and ZNF665. (C) Zinc fingerprints of ZNF160 and ZNF665 with the scaled minor allele frequency (MAF) of identified missense variants indicated on the sides. Grey lines indicate identical zinc fingerprints (D) Consensus DNA binding motifs of ZNF160 and ZNF665. (E) Venn diagram of ChIP-exo peaks of ZNF160 and ZNF665 in 293T cells.

### The lexicon of human KZFP genomic targets is strongly biased towards TEs

We previously identified the genomic targets of 242 human KZFPs through chromatin immunoprecipitation followed by DNA sequencing (ChIP-Seq) (Imbeault et al., 2017; Helleboid et al., 2019). We extended these analyses to an additional 110 family members, similarly using HA-tagged derivatives overexpressed in 293T cells transduced with dox-inducible lentiviral vectors (Imbeault et al., 2017). Of the remaining 26 KZFPs, DNA could not be successfully synthesized in 3 cases, while transduction yielded no or little protein in another 23 (Figure 4 and Table S2). Peak numbers varied between experiments, but we observed that this parameter did not reflect the quality of the ChIP-Seq as very specific DNA sequences were enriched for KZFPs with low peak numbers. Accordingly, after removing sequences present in all ChIPs irrespective of the target protein, we considered enrichment over specific DNA sequences as the main criterion for assessing the quality of an experiment. Integrating our cumulated data with those previously obtained by other groups (Frietze et al. 2010; ENCODE Project Consortium 2012; Yan et al. 2013; Schmitges et al. 2016; Venkataraman et al. 2018; Haring et al. 2021; Partridge et al. 2020), we could build a lexicon constituted by the genomic targets of 358 human KZFPs, including replicates for 78 of them (Table S2).

**Figure 4:**
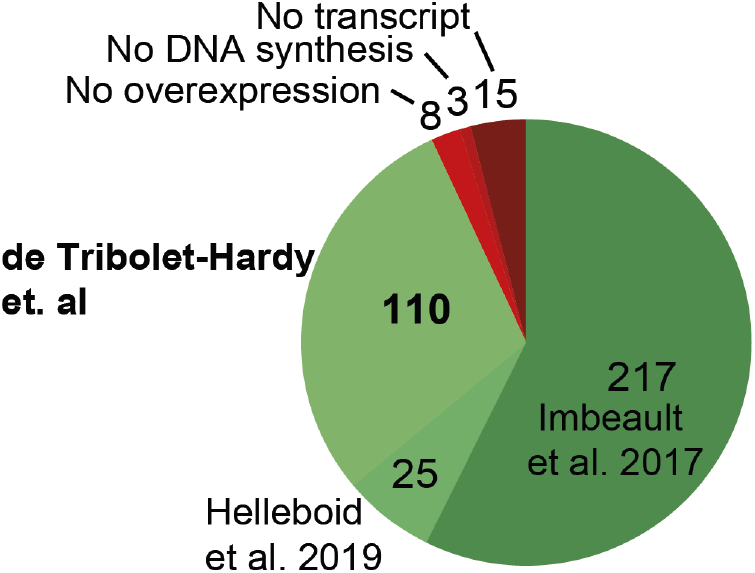
Profiling of the human KZFPs. A) Pie-chart of the data on all 378 protein-coding KZFPs. “No overexpression” indicates the number of KZFPs where the codon-optimized construct did not yield sufficient protein. “No transcripts” represents KZFPs with no annotated transcript containing both the KRAB and zinc finger domains simultaneously. “No DNA synthesis” indicates the number of KZFPs CDS that could not be synthesized, with a minimum of two tries.

As previously observed through studies on smaller subsets of KZFPs (Imbeault et al. 2017; Najafabadi et al. 2015; Helleboid et al. 2019), our integrated analysis confirmed that the vast majority of human KZFPs are foremost enriched at TE-derived loci (Figure 5A and S4). Due to the close relationship between different TE subfamilies and to the production of enrichments rather than binary results by the ChIP-seq technique, we generally have several TE subfamilies significantly enriched in a single experiment. However, in most cases a few subfamilies stand out as being much more enriched than others. We defined the identified sequences as primary targets of the ChIP’ed KZFPs if within an arbitrary cut-off (10% of the log10 of the lowest False Detection Rate -FDR-) and designated the remainder as potential secondary targets (Table S3 and Figure S5). Of 349 KZFPs with clearly identifiable targets, 120 were preferentially enriched at ERVs, 70 at LINES, 21 at SINEs/Alus, 32 significantly bound SVAs and 11 were rather found at DNA transposons. The remaining 71 KZFPs mapped to a mixture of low complexity or simple repeats, satellite DNAs and tRNAs. For 24, no particular class of genomic entity could be singled out perhaps due in part to difficulties in sequences alignment, notably in telomeric and centromeric regions (Figure 5A and S5A with further details on https://tronoapps.epfl.ch/web/krabopedia/). Altogether, about two-thirds of know human TE subfamilies are primary targets of at least one KZFP, this number exceeding 95% if secondary targets are also considered (Figure 5B). Thus, both a large majority of KZFPs bind TEs and a large majority of TEs are bound by KZFPs, further strengthening the evolutionary and functional link between these two genetic entities. Confirming with this full dataset a trend noted previously (Imbeault et al., 2017), the evolutionary times of TEs and of their controlling KZFPs most often coincided (Figure 5C). When comparing the age of KZFPs (red line) with the age of their targets (black bars), contemporary waves of KZFPs and TEs emergence can be observed. For example, LINE1 subsets and ERVL that emerged some 105 million years ago are predominantly bound by KZFPs of similar ages, an observation that holds true for the younger ERV1 and ERVK and their cognate ligands (see Table S4). However, such evolutionary pairing does not apply to SINE and SVA elements, both of which are targeted primarily by older KZFPs, including family members not interacting with KAP1 (red dots). Another interesting case are LINE2 elements, half of which are targeted by contemporary KZFPs whereas the other half is bound by evolutionarily younger ligands.

**Figure 5:**
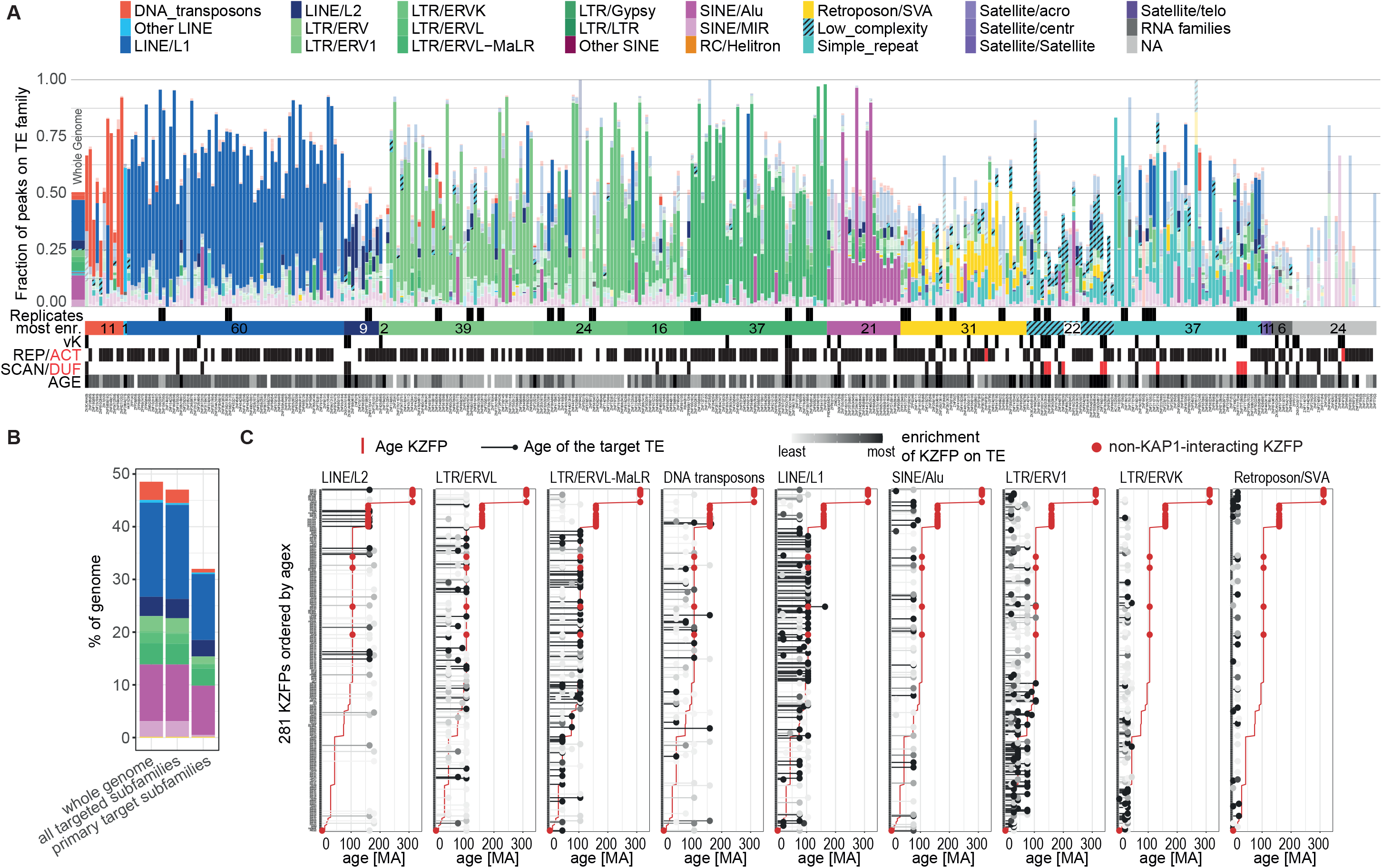
Targets of human KZFPs. A) Bar graph showing the fraction of peaks over repetitive element (RE) families for all our conducted experiments (x-axis) (external data is shown in Figure S5A), ordered by the most enriched family, indicated by the horizontal bar below along with the number of KZFPs for each category. Significant enrichments (FDR < 0.05) are shown in fully opaque colours whereas non-significant enrichments are transparent. The leftmost bar shows the percentage of the genome occupied by each RE family. Replicate experiments are indicated by black squares above the horizontal bar. Different aspects of each KZFP are shown below the horizontal bar: vK=variant KRAB according to (Helleboid et al. 2019), REP/ACT= repressor and activator KZFPs according to (Tycko et al. 2020), SCAN/DUF= KZFP carrying an additional SCAN or DUF3669 domain. Age: Black= >105 myo, dark grey= <105 myo years (placental mammals), light grey= < 74 myo years (primates), white = No data. The total number of peaks per experiment is indicated in brackets after the KZFP name below each bar. B) Bar graph showing the genome occupancy of targeted TE subfamilies. The left stack of bars shows the fractions of the genome covered by TEs, the central stack shows the coverage by all TE subfamilies which are targeted by a KZFPs (FDR < 0.05) and the right stack shows the coverage of the TE subfamilies which are the primary target of one or more KZFPs (10% highest -log10[FDR]). Bars are coloured according to the TE families the subfamilies belong to, with the same colour code as in panel A. C) Age of KZFP and their target TEs. KZFPs (rows) are ordered by age, shown as a red line. Their targets are split into different subplots by family (excluding families targeted by less than 20 KZFPs) and their age is shown as black or grey bars with a dot on top. The grey level of the TE targets shows the level of enrichment of the given KZFP for the subfamily with black showing the target with the highest -log10(FDR) linearly scaling to 0 (white). If the KZFP is enriched on several subfamilies of the same family the lowest FDR is shown. Red dots indicate KZFPs which are unlikely to interact with TRIM28 as defined by Helleboid et al. 2019.

### TEs are avid KZFP recruiters

Our data confirm that most TE subfamilies are bound by more than one KZFP (Figures 6A and S6A). This feature is particularly striking for SVAs, considering the relatively young age of this class of retrotransposons (around 15 million years for the oldest SVA-A), their small size (on average 2,500 bp) and the low number of their integrants (some 3,500 for the entire family) (Kojima 2018; H. Wang et al. 2005). SVAs are non-autonomous composite elements made up by the juxtaposition of an Alu-like sequence, a Variable Number of Tandem Repeats (VNTR) and an ERV-derived 3’ region called SINE-R, somewhat of a misnomer since not related to the SINE family of TEs (Ono et al. 1987). A few tens of SVAs (belonging to the youngest, human-specific SVA-F subset) are still transposition competent, using for their replication the reverse transcriptase and endonuclease activities provided by LINE1 *in trans.* Many SVAs, notably from the SVA-D subgroup, provide enhancers active in early embryogenesis and/or in adult tissues, including at KZFP genes clusters (Haring et al. 2021; Pontis et al. 2019, 2022; Gianfrancesco et al. 2019). The distribution of KZFP binding sites over SVA sequences reveals three distinct patterns (Figure 6B). ZNF705A, B, D and E as well as ZNF282 and ZNF780A are enriched over the Alu-like segment, whereas the previously described (Jacobs et al. 2014) ZNF611 and ZNF91 bind to the 5’ end of the VNTR. However, the vast majority of ChIP signals overlap with the more distal, highly variable part of the VNTR, where 29 KZFPs are enriched.. However, ZNF141, which yielded the strongest signal (Figure 6C), was also enriched over L1PA3 and L1PA2 integrants and SATR1 Satellite repeats (Figure S6B), and the same previously identified binding motif (Weirauch et al. 2014) was found in all three targets, confirming *bona fide* binding (Figure S6C). LINE1 integrants, at least when full-length, are also targeted by multiple KZFPs, some binding towards the 5’ end of these integrants repressing their transcription and others recognizing downstream regions dispersed all the way to their 3’ end (Figure 6D). Furthermore, comparing the KZFP recruitment patterns of recent and older LINE1 subfamilies reveals differential evolutionary forces at play. For instance, the KAP1-binding ZNF93 repressor, which emerged in the last common ancestor of apes and Old-World monkeys, recognizes the promoter regions of ~27 myo L1PA6 to ~16 myo L1PA3 but not that of ~3 myo L1Hs (Jacobs et al. 2014), whereas the more distally binding ZNF382 and ZNF490, both also KAP1 recruiters but ~105 myo (Helleboid et al., 2019), are binding integrants from all these LINE1 subsets.

**Figure 6:**
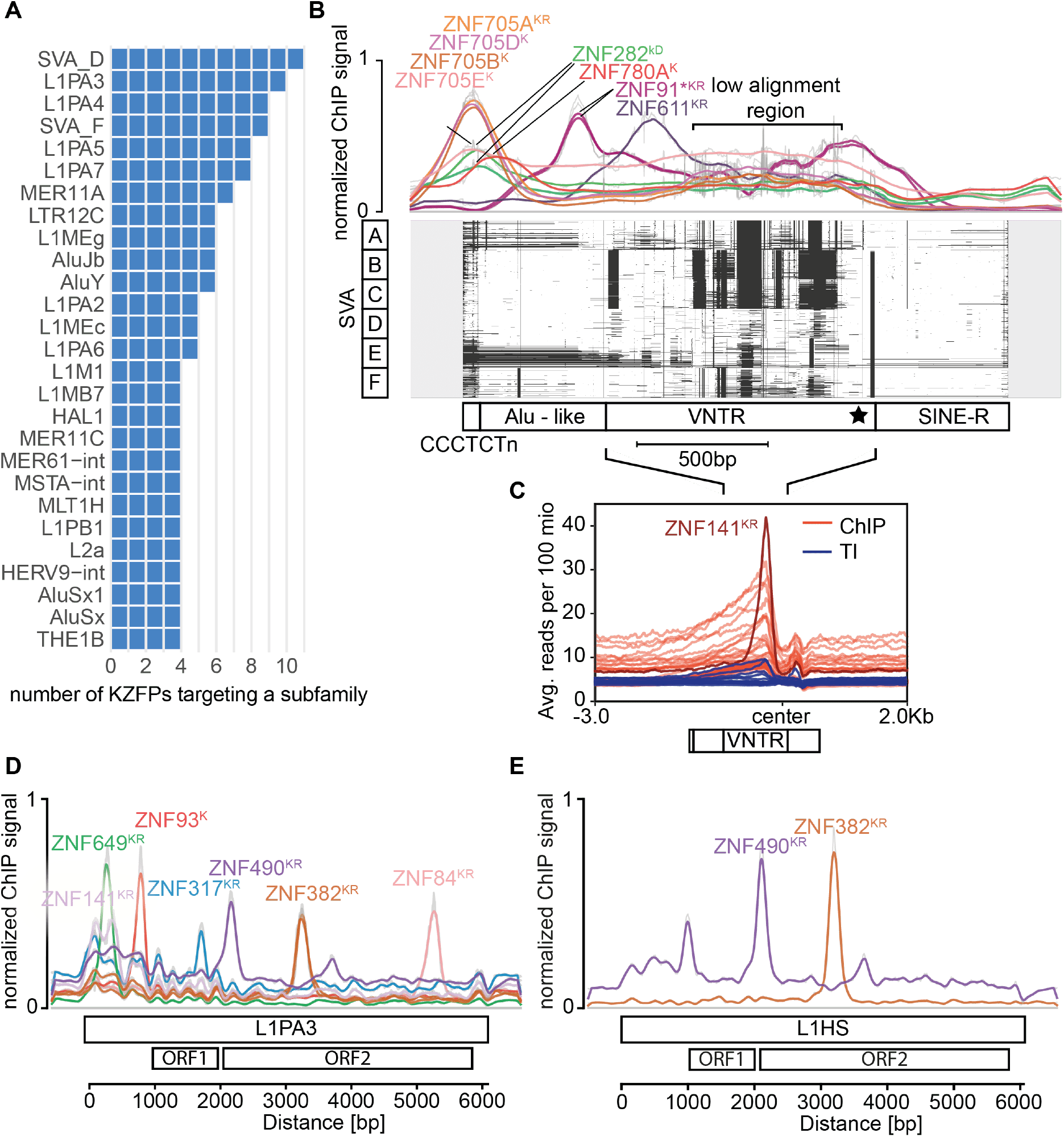
TE families are targeted by multiple KZFPs. A) Bar graphs showing the TE subfamilies targeted by the largest number of different KZFPs. Only KZFPs targeting the subfamily as their primary targets where considered (-log10(FDR) within 10% of the highest -log10(FDR) for that KZFP). B) KZFP signal over the multiple sequence alignment (MSA) of SVA subfamilies A to F. Top: Line graph of the normalized cumulative reads for each position from the indicated ChIP-seq and -exo experiments. External datasets are marked with stars. Bottom: MSA plot of 100 of the longest SVA sequences for each subfamily indicated on the left, 200 bp of non-aligned extensions are added around elements shown in grey, white depicts aligned regions and black gaps in the alignment. For visibility, places in the alignment (columns) with more than 85% gaps were removed. The approximate different domains of the SVAs are indicated below, adapted from (Hancks and Kazazian 2010), the star indicates the centre region for panel C. C) Signal over the low alignment region of the remaining SVA binders centred on the 3’ end of the VNTR (without alignment of sequences). ChIP signals for KZFPS enriched on SVAs are shown in red (ZFP57, ZFP92, ZNF14, ZNF141, ZNF155, ZNF215, ZNF25, ZNF256, ZNF263, ZNF268, ZNF28, ZNF30, ZNF41, ZNF415, ZNF461, ZNF500, ZNF556, ZNF560, ZNF57, ZNF587B, ZNF597, ZNF624, ZNF641, ZNF689, ZNF699, ZNF747, ZNF813, ZNF852 and ZNF878) with the signal for ZNF141 is shown in dark red. Input signals for the presented ChIPs are shown in blue. Multi-mapped reads where included for the signals in B and C. D and E) Binding sites of KZFPs on L1PA3 and L1HS elements. Elements were aligned the way as in A and the normalized ChIP-seq and -exo signals are shown for each aligned position. External datasets are marked with stars. K=Standard KRAB, k= variant KRAB, D=DUF domain, R=Repressor; according to (Helleboid et al. 2019; Tycko et al. 2020). D) 1000 L1PA3 elements were aligned. E) 382 full length L1HS elements were aligned.

### Multi-pronged modes of evolution of the TE-KZFP interaction

KZFPs present within the same cluster are often related in sequence since in many cases derived from each other (Lukic et al. 2014). For example, the ~105 myo ZFP69 and ZFP69B, encoded side-by-side in chr1.1 cluster (Figure 7A), display significant similarities in their zinc fingerprints and DNA binding motives (Figure 7B and C). However, their primary targets differ, with ZFP69 preferentially recognizing a mammalian-specific LINE1 element and ZNF69B favouring LTR HERVH-int (Figure 7D and E). Yet, an examination of their secondary targets identifies L1MC1 at a significant frequency in both cases, suggesting that the ancestral ZFP69 might have targeted this element. Other examples of KZFPs encoded by neighboring genes and displaying distinct primary but shared secondary targets include ZNF695, ZNF669 and ZNF124, transcribed from a chr1 *KZFP* gene cluster, and ZNF354A, ZNF354B, ZNF454, ZNF879 and ZNF354C, encoded next to each other on chr5 (Figure S7A).

**Figure 7:**
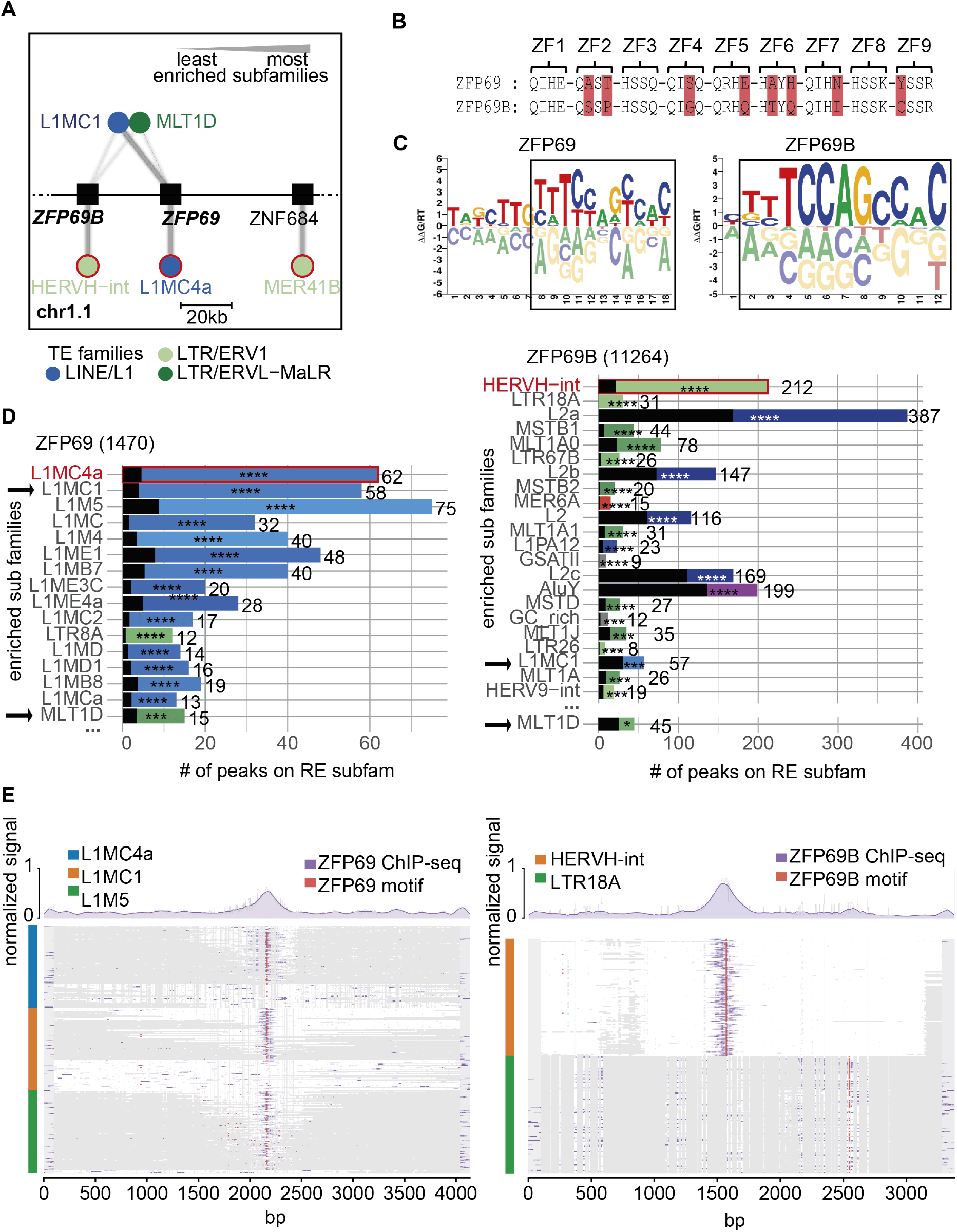
Evolution of TE-KZFP interaction. A) Network for cluster chr1.1 where targets (circles) of each KZFP (squares) are shown as connected edges and the amount of binding is represented by the line thickness. The thickest line for each KZFP represents the TE subfamily with the highest -log10(FDR) and then scales linearly to the lowest value. For visibility, only the best targets (below) and shared targets (above) are shown. The TE subfamilies are coloured according to their families. Primary targets for each KZFP are highlighted in red. B) Zinc fingerprints of ZFP69 and ZFP69B. The DNA-contacting amino acids for zinc finger (ZF) are shown, differences are highlighted in red. C) DNA binding motifs of ZFP69 and ZFP69B as identified by (Weirauch et al. 2014). Regions of high similarity are framed by a black square D) Enrichment of peaks over different repetitive element subfamilies. Subfamilies with FDR < 0.01 are shown. The width of the coloured bars represents the number of peaks per subfamily also shown as a number on the right of the bar. The black transparent bars represent the expected number of peaks following a random distribution. The FDR of the enrichment is shown with stars (FDR < 0.0001 = ****, < 0.001=***, < 0.01=**, < 0.05=*, >= 0.05 = n.s). Rows are ordered by FDR. The number next to the title indicates the total number of peaks for the experiment. Primary targets for each KZFP are highlighted in red and shared subfamilies between the two panels are indicated by black arrows. E) Multiple sequence alignment (MSA) over the 3 most enriched targets of ZFP69 (left) and ZFP69B (right). Up to 200 elements for the indicated targets (blue, orange and green) where aligned, selecting first elements overlapping with a peak and then the longest elements. White regions in the plots indicate aligned sequences, grey regions indicate gaps. The signal of ZFP69 and ZPF96B ChIPs was laid over their respective alignments in purple. The location of their motifs from panel C are shown in red. The normalized signal can be seen as a line plot above the MSA plot.

The recruitment of multiple KZFPs by given TE integrants and strong evidence for the role of differential selective pressures (e.g. the arms race model for ZNF93 and the transition between L1PA3 and L1PA2 vs. the consistent recruitment of ZNF84 and ZNF282 by multiple generations of LINE1) suggest a multimodal evolution of the TE-KZFP relationship. This is supported by the finding that KZFPs binding to a same TE subgroup are commonly encoded in different KZFP gene clusters, as exemplified with MER11A, L1PA3 and SVAs (Figure 8A-C), and in a somewhat linear relationship between the number of KZFPs and corresponding source gene clusters involved in recognizing a given element, with an approximate ratio KZFP/source cluster of 2:1, down to 1:1 when considering only primary targets (Figure 8D).

**Figure 8:**
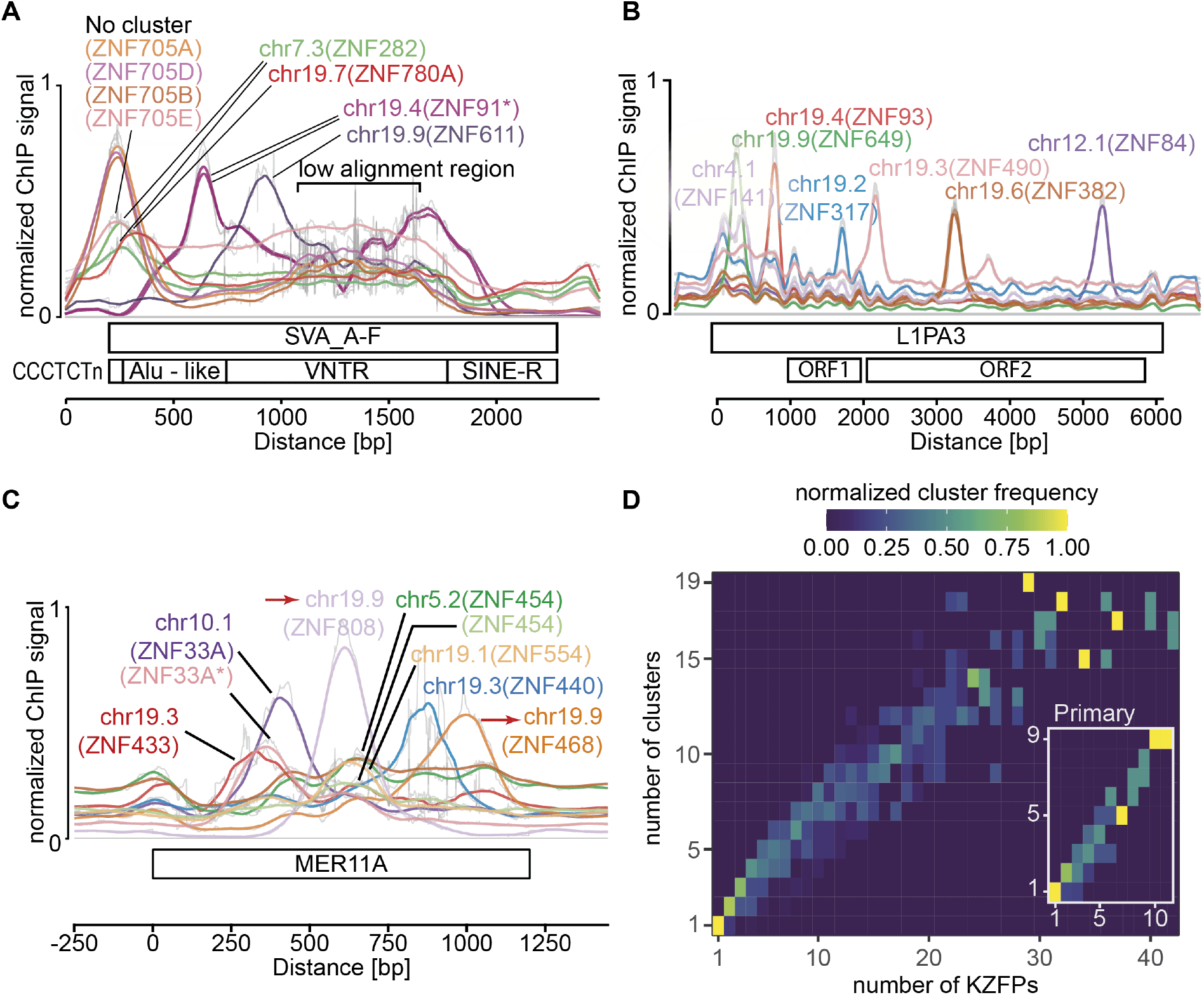
Localization of multiple KZFPs targeting the same TE subfamily. A-C) Cluster location of the indicated KZFPs, which primarily bind the respective TE subfamilies. Duplicated clusters are marked with red arrows, external datasets are marked with stars. A) Alignment of up 200 of the longest SVA_A to SVA_F elements B)-C) Alignment of 1000 L1PA3 and MER11A elements. D) Heatmap showing the number of KZFP-clusters (y-axis) the KZFPs targeting the same TE subfamily (x-axis) are localized in. Colors show the normalized distribution of the number of clusters for each number of KZFPs. Large panel: All KZFPs targeting a subfamily (FDR < 0.05) are considered. Sub panel: Only KZFPs primarily targeting a subfamily (-log10(FDR) within 10% of the highest -log10(FDR)) are considered.

## DISCUSSION

This extensive mapping of human KZFPs genomic targets confirms that in their vast majority these proteins recognize sequences embedded in transposable elements. Altogether, genomic binding sites have now been characterized for 358 out of 378 family members, revealing that 254 of them have a TE as their primary target. Conversely, our results indicate that most TEs can be recognized by a KZFP, and many by more than one. Since only ~1:1000 TE is still capable of transposition, it lends strong credence to our early proposal (Trono 2015) that the evolutionary selection and maintenance of KZFP genes has been geared towards the domestication of TE-embedded regulatory sequences (TEeRS) rather than driven by the need to block the spread of these genetic invaders, even though the two are not mutually exclusive. For instance, ZNF93 is the prototype of a KZFP initially involved in some sort of TE-host arm race, the fixation of which coincided with the emergence of the L1PA6 generation of LINE1 before their L1PA3 descendants escaped its control by deleting its binding site, some 15 to 20 million years later (Jacobs et al. 2014). Yet, even for ZNF93, control of transposition appears to have been only a temporary function, as this KZFP keeps recruiting a KAP1-associated repressor complex to the 5’ end of thousands of L1PA6 to L1PA3 integrants, all of which became transposition-defective millions of years ago.

Most remarkable is the ability by many integrants from the LINE, ERV or SVA families to recruit over different sites in their sequences multiple KZFPs, most bearing a standard KRAB domain with verified repressor potential. While for ERVs and SVAs these KZFP binding sites are generally clustered close to known promoter or enhancer elements, with for instance a concentration of KZFP-recruiting motifs within ERV LTRs, the distribution of KZFP peaks over the whole sequence of LINE1 integrants, including in the central and 3’ regions, is intriguing. First, it confirms that transposition-deficient TEs are major genomic docking sites for KZFPs, since many LINE1 integrants are 5’ deleted, hence devoid of 5’ promoter, due to incomplete retrotranscription. Second, it indicates that these distally situated LINE1 sequences and their KZFP ligands must accomplish some biological functions, the nature of which is still largely to decipher. Third, it calls for studies examining the spatio-temporal regulation and biological impact of the recruitment of these KZFPs on their TE targets both alone and, when relevant, in combinations.

The evolutionary conservation of individual KZFPs correlates with the genetic constraint imposed on their coding sequences: older KZFP genes display lower degrees of genetic variation in the human population than their more recent counterparts, notably at positions encoding amino acids predicted to dictate the DNA binding specificity of their products. Yet the target sequences of these highly conserved KZFPs reveal an interesting dichotomy, encompassing both very old TEs such as L2 or DNA transposons and evolutionary recent elements such as SVAs or ERV1s. Furthermore, many of these KZFPs harbor variant KRAB domains that do not interact with KAP1 and associated epigenetic modifiers but with other types of protein complexes, and are devoid of repressor activity (Helleboid et al. 2019; Tycko et al. 2020). This strongly supports a model whereby TEs serve as vectors of *cis*-acting regulatory sequences of a broad functional diversity.

In contrast to older family members, evolutionarily recent KZFPs almost universally target TEs, often have paralogs and display a KAP1-centered protein interactome primarily consistent with transcriptional repression (Helleboid et al. 2019; Tycko et al. 2020). The greater degree of polymorphism observed in the human population at positions determining the genomic targets of these recently emerged KZFPs may be explained by the absence of TEs forcing fixation of at least part of their ZF-coding sequences. Interestingly, differentials in coding constraint varied within unequivocally identified sets of KZFP paralogs, being very narrow in some cases (e.g., *ZNF75A* and *ZNF75D; ZFP30* and *ZFP82*) and quite broad in others (e.g., *ZNF160* and *ZNF665; ZNF181* and *ZNF302*). No single parameter could account for these differences. For instance, ZNF75A and ZNF75D both recognize the 3’ end of KZFP genes, while ZFP30 and ZFP82 respectively bind LINEs and SINEs, that is, completely distinct sets of genomic targets. As well, ZNF679-ZNF716 and ZNF600-ZNF611 are two pairs of evolutionarily recent (<20 myo) paralogs, yet they present with coding constraint differentials that are negligible for the former and pronounced for the latter. Still, it is noteworthy that for paralogs of detectably distinct ages, the older gene is generally more constrained than its duplication product, recapitulating a trend noted for the entire KZFP family.

The KZFP gene pool of a lineage undergoes a high evolutionary turnover, as indicated by the mammalian- and primate-specificity of 88% and 32%, respectively, of human family members. The gene duplication mechanism underlying this phenomenon allows for an efficient diversification of the *trans*-regulatory space without losing track of physiology, as new TEeRS emerging by genetic drift of the host TE pool can be controlled and potentially exploited without unleashing the perturbation potential of older TEeRS. This smooth transition model is supported by an examination of the secondary targets of paralogs such as ZFP69 and ZFP69B, which suggests that an ancestral *ZFP69* gene targeting L1MC elements duplicated to have the original gene conserve its affinity for this TE and the zinc fingerprint of its copy drift to become fixed upon recognition of a later emerged HERV. In this system, even if only a fraction of newcomer genes ends up positively selected, a rapid flux of new candidates, on both the TE and KZFP sides, fuels the evolution of a lineage’s regulome. Most frequently, due to environmental and physiological constraints, this will result in purely mechanistic speciation, with conservation of biological processes but turnover of some of their *cis-* (the TEeRS) and trans- (the KZFPs) regulators, as during early embryogenesis or gametogenesis (Pontis et al. 2022, 2019; Barnada et al. 2022; Xiang et al. 2022). Occasionally however, it may give raise to new traits, notably in organ systems where the range of phenotypes compatible with reproductive life hence trans-generational inheritance is greater, as suggested by the increasingly recognized importance of TE/KZFP-mediated regulation in the developing human brain (Turelli et al. 2020; Playfoot et al. 2021, 2022; Johansson et al. 2022; Patoori et al. 2022; Farmiloe et al. 2020; Nowick et al. 2009).

## METHODS

### Census of the human KRAB Zinc Finger protein clusters

KZFP pairs were detected and their age defined as described in (Imbeault et al. 2017). In short, the human genome (hg19) was translated in 6 reading frames and scanned for zinc finger and KRAB domains using Hidden-Markov-Models (Pfam (El-Gebali et al. 2019): KRAB (PF01352) and zf-C2H2 (PF00096)). Hits for KRAB and zinc finger domains were combined based on proximity and strandness and then manually curated and integrated with existing gene or pseudogene annotations. Their age is based on sequencing similarity with orthologues in other species. The KZFP clusters were defined as having at least 3 KZFPs that are no more than 250 kb apart from the centre of another member, consistent with (Huntley et al. 2006). The clusters are named after their chromosome and then numbered starting from the short arm of the chromosome. The size of chromosomes and positions of centromeres were taken from UCSC genome browser annotation data for hg19 (Haeussler et al. 2019).

### Primate phylogeny and natural selection

The time of divergence (i.e. branch lengths) between human, chimpanzee, gorilla, orangutan, macaque, marmoset, tarsier, galago (a.k.a. bush baby) and mouse lemur was obtained from 10KTrees, which uses Bayesian inference to estimate these (Arnold et al. 2010). Measures of natural selection in terms of dN/dS across the nine primate species listed above was obtained with PAML (v4.4) as previously described (McLaren et al. 2015).

### Human genetic variation data

Human genetic exome and whole genome sequencing data were obtained from The Genome Aggregation Database (gnomAD) (Lek et al. 2016; Karczewski et al. 2020) (release-2.0.2) for 123,136 and 15,496 individuals, respectively. The released genetic data was processed and filtered through several steps to guarantee that only high-quality variants were included. First, all variants +/- 1kb around the KZFP canonical transcripts as defined by Ensembl (v75, hg19) were extracted and filtered for variant quality, thus only retaining variants annotated as “PASS”.

Second, all indels were normalized and multiallelic variants split using BCFTOOLS (v1.8) and reannotated with the Variant Effect Predictor (McLaren et al. 2016) and LOFTEE (v0.3beta). Third, all missense and loss of function (LoF) variants, defined as either frameshift, stop-gain or splice variants, were extracted from both the exome and whole genome datasets and either low confidence or flagged LoF variants were removed. The latter was primarily due to LoF variants found in the last 5% of the canonical transcript. Since genomic sequencing methods can yield variable coverage of genetic regions, especially when it comes to exome sequencing that is dependent on the capture of previously annotated protein-coding genes, we excluded all canonical transcripts having an average per-base coverage < 20x. Thus, bringing the total number of included KZFPs to 361. Furthermore, exons with an average per-base coverage < 20x were also removed, and the lengths of the coding sequences used later for normalizations were adjusted accordingly. Finally, the filtered exome and genome datasets were combined, and the allele counts and frequencies for all variants were recalculated, prior to the removal of all singletons (allele count = 1) to hinder inflation of observed mutational events due to potential technical artefacts.

### Domain and site specifications

The genomic positions of the C2H2 zinc finger domains were obtained from the Ensembl database (v75, hg19). For each KZFP, only the ones from the canonical transcripts (as defined by Ensembl) were considered. The positions of the specific amino acids within these domains were computationally annotated. z-scores for the cysteine and histidine (C2H2) residues were calculated with the number of missense variants normalized to the number of zinc finger domains within the canonical transcript of each KZFP. For missense and LoF variants spanning either the whole CDS or a full protein domain, the number of variants per gene, x, was normalized by the length of the canonical coding sequence prior to z-score transformations.

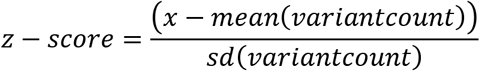

### Cell Lines

HEK293T cells overexpressing HA-tagged KZFPs were generated as described in Imbeault et al., 2017. In short, cDNAs from the human KZFPs were codon-optimized and synthesized using the GeneArt service from ThermoFisher (former Life Technologies). Sequences were cloned into the doxycycline inducible expression vector pTRE-3HA which yields a C-terminally tagged proteins. Stable cell lines were generated using Lentivector transduction of mycoplasma free HEK293T cells as described on *http://tronolab.epfl.ch.* Presence and integrity of the integrated plasmids were verified using Sanger sequencing (primers: CMV1f: GGAGGCCTATATAAGCAGAGCTCGT, PGK4b: CGAACGGACGTGAAGAATGTGCGAGA) and KZFP expression was verified via western blot with an anti-HA antibody (ref. 12013819001, Roche) after >48h induction with 1ng/ml doxycycline. HEK293T cells were chosen in order to have a consistent cell line and genomic background for all conducted experiments.

### ChIP-seq

Chromatin was prepared as described in Imbeault et al., 2017 and ChIP-seq was performed as described in Iouranova et al., 2022. In short: 30 mio KZFP expressing HEK293T cells were used after induction for more than 48h with 1ng/ml doxycycline. Cells were crosslinked with 1% methanol free formaldehyde for 10min before nuclear extraction followed by sonication in a Covaris E220 sonicator resulting in DNA fragments between 200-500bp. IP was performed overnight using 15μg anti-HA.11 antibody (BioLegend ref: 901503) coupled to 75ul Dynabeads Protein G (Invitrogen ref: 10009D). 10 ng of material for both total inputs and chromatin immunoprecipitated samples were used for library preparation. After end-repair and A-tailing, Illumina IDT indexes were ligated to the samples. Aliquots were tested in qPCR to determine the optimal number of PCR cycles needed to amplify each library without reaching saturation. Libraries were size-selected using Ampure XP beads (Beckman Coulter), quality-checked on a Bioanalyzer DNA high sensitivity chip (Agilent) and quantified with a Qubit dsDNA HS assay kit (Qubit 2.0 Fluorometer, Invitrogen) using Illumina adapters. Libraries were sequenced as indicated on GEO: GSE200964.

### Processing of ChIP-seq and ChIP-exo data

Both previously published (Helleboid et al. 2019; Imbeault et al. 2017) and new data were processed together. Reads were mapped to the human genome assembly hg19 using Bowtie2 short read aligner v2.3.5.1 (Langmead and Salzberg 2012), using the --sensitive-local parameter. Prior to peak calling multi-mapped reads (MAPQ < 10), blacklisted regions and regions with high levels in input samples (greylist) defined by the R package GreyListChIP (Brown 2022) were removed. Performing the analyses including multi-mapped reads slightly changed the set of peaks, adding some and removing others, but did not influence the vast majority of target enrichments (data not shown). Peaks were called using MACS2 v2.2.4 (Zhang et al. 2008) with defaults parameters except for *-q 0.01* and *--keep-dup all.*

### External ChIP-seq data

To find KZFP ChIP-seqs done by others the programmatic access to GEO (Barrett et al. 2013) eSearch and eFetch functions were used to search and retrieve submissions containing any KZFP name but not the keywords “RNA” or “H3K”. The resulting hits were then manually curated using the GEOquery R package (Davis 2022) in order to get bed files from ChIP-seq experiments. Peaks not called on hg19 were lifted over to hg19 using liftover from rtracklayer (Lawrence et al. 2022) and chain files from UCSC (Haeussler et al. 2019).

### Enrichment on repeats

Repeat enrichment analyses from ChIP data were performed using pyTEnrich (https://alexdray86.github.io/pyTEnrich). Repeat annotations were obtained from UCSC (http://hgdownload.soe.ucsc.edu/goldenPath/hg19/database/rmsk.txt.gz). The genome subset for the enrichment was generated by identifying regions with 0 coverage for each ChIP-seq and ChIP-exo using bedtools genomcov, merging regions with less than 100 bp distance and overlapping the resulting files with bedtools intersect. The resulting intervals were then filtered to be bigger than 40kb and removed from the analysis along with the Y-chromosome which is absent in HEK293T cells. Enrichments with FDR < 0.05 are considered significant. In order to normalize FDR between experiments -log10(FDR) were divided by their maximum yielding a scale from 0 to 1 or least to most enriched, values above 0.9 on this scale are considered primary targets.

### Multiple sequence alignment plot and line plots

Multiple sequence alignment (MSA) plots where made as described in Iouranova et al., 2022. In short: FASTA sequences for the indicated subfamilies were extracted from the hg19 genome assembly, aligned using individually using MAFFT (Katoh and Standley 2013) with parameters --reorder –auto, and then merged together using MAFFT’s -merge option. To increase readability, positions in the alignment (columns) with more than 85% gaps were removed. To capture signal at the border the alignments are extended by 200-500bp of unaligned sequences. ChIP-seq and -exo signals are scaled for each line (row) to the [0,1] interval before being superimposed on the alignments. Average ChIP-seq signals across all rows are plotted on top of the alignments or without alignment for Figures 2 and 5. Motives were taken from Cisbp (Weirauch et al. 2014) converted to position weight matrixes and scanned for in the human genome (hg19) using PWMscan (Ambrosini et al. 2018) with default settings. Line plots in Figure3B were generated using deeptools plotProfile (Ramírez et al. 2016). SVA for all subfamilies (A-F) were centred on a well conserved region on the edge of the VNTR with the consensus sequence ACTAAGAAAAATTCTTCTGCCTTGGG.

## DATA ACCESS

ChIP-seq and -exo data have been deposited in the Gene Expression Omnibus (GEO) database with the accession number GSE200964. Cumulated information on human KZFPs is available in our KRABopedia database at https://tronoapps.epfl.ch/web/krabopedia/.

## ACKNOWLEDGMENTS

We thank Charlène Raclot and Kerim Benbouhafs for technical assistance, Bastien Mangeat and the Gene Expression Core Facility at EPFL for sequencing and SCITAS for computing infrastructure.

## REFERENCES

Ambrosini G, Groux R, Bucher P. 2018. PWMScan: a fast tool for scanning entire genomes with a position-specific weight matrix. Bioinformatics 34: 2483–2484.

Arnold C, Matthews LJ, Nunn CL. 2010. The 10kTrees website: A new online resource for primate phylogeny. Evol Anthropol Issues News Rev 19: 114–118.

Barnada SM, Isopi A, Tejada-Martinez D, Goubert C, Patoori S, Pagliaroli L, Tracewell M, Trizzino M. 2022. Genomic features underlie the co-option of SVA transposons as cis-regulatory elements in human pluripotent stem cells. 2022.01.10.475682. https://www.biorxiv.org/content/10.1101/2022.01.10.475682v1 (Accessed April 14, 2022).

Barrett T, Wilhite SE, Ledoux P, Evangelista C, Kim IF, Tomashevsky M, Marshall KA, Phillippy KH, Sherman PM, Holko M, et al. 2013. NCBI GEO: archive for functional genomics data sets—update. Nucleic Acids Res 41: D991–D995.

Britten RJ, Davidson EH. 1969. Gene Regulation for Higher Cells: A Theory. Science 165: 349–357.

Brown G. 2022. GreyListChIP: Grey Lists -- Mask Artefact Regions Based on ChIP Inputs. https://bioconductor.org/packages/GreyListChIP/.

Bruno M, Mahgoub M, Macfarlan TS. 2019. The Arms Race Between KRAB-Zinc Finger Proteins and Endogenous Retroelements and Its Impact on Mammals. Annu Rev Genet 53: 393–416.

Chen W, Schwalie PC, Pankevich EV, Gubelmann C, Raghav SK, Dainese R, Cassano M, Imbeault M, Jang SM, Russeil J, et al. 2019. ZFP30 promotes adipogenesis through the KAP1-mediated activation of a retrotransposon-derived Pparg2 enhancer. Nat Commun 10: 1809.

Davis S. 2022. GEOquery: Get data from NCBI Gene Expression Omnibus (GEO). https://bioconductor.org/packages/GEOquery/.

Durnaoglu S, Lee S-K, Ahnn J. 2021. Human Endogenous Retroviruses as Gene Expression Regulators: Insights from Animal Models into Human Diseases. Mol Cells 44: 861–878.

El-Gebali S, Mistry J, Bateman A, Eddy SR, Luciani A, Potter SC, Qureshi M, Richardson LJ, Salazar GA, Smart A, et al. 2019. The Pfam protein families database in 2019. Nucleic Acids Res 47: D427–D432.

Emerson RO, Thomas JH. 2009. Adaptive Evolution in Zinc Finger Transcription Factors. PLOS Genet 5: e1000325.

ENCODE Project Consortium. 2012. An integrated encyclopedia of DNA elements in the human genome. Nature 489: 57–74.

Farmiloe G, Lodewijk GA, Robben SF, van Bree EJ, Jacobs FMJ. 2020. Widespread correlation of KRAB zinc finger protein binding with brain-developmental gene expression patterns. Philos Trans R Soc B Biol Sci 375: 20190333.

Frietze S, Lan X, Jin VX, Farnham PJ. 2010. Genomic targets of the KRAB and SCAN domain-containing zinc finger protein 263. J Biol Chem 285: 1393–1403.

Gianfrancesco O, Geary B, Savage AL, Billingsley KJ, Bubb VJ, Quinn JP. 2019. The Role of SINE-VNTR-Alu (SVA) Retrotransposons in Shaping the Human Genome. Int J Mol Sci 20: E5977.

GTEx Consortium, Aguet F, Brown AA, Castel SE, Davis JR, He Y, Jo B, Mohammadi P, Park Y, Parsana P, et al. 2017. Genetic effects on gene expression across human tissues. Nature 550: 204.

Haeussler M, Zweig AS, Tyner C, Speir ML, Rosenbloom KR, Raney BJ, Lee CM, Lee BT, Hinrichs AS, Gonzalez JN, et al. 2019. The UCSC Genome Browser database: 2019 update. Nucleic Acids Res 47: D853–D858.

Hancks DC, Kazazian H. 2010. SVA retrotransposons: Evolution and genetic instability. Semin Cancer Biol 20: 234–245.

Hancks DC, Kazazian HH. 2016. Roles for retrotransposon insertions in human disease. Mob DNA 7: 9.

Haring NL, van Bree EJ, Jordaan WS, Roels JRE, Sotomayor GC, Hey TM, White FTG, Galland MD, Smidt MP, Jacobs FMJ. 2021. ZNF91 deletion in human embryonic stem cells leads to ectopic activation of SVA retrotransposons and up-regulation of KRAB zinc finger gene clusters. Genome Res 31: 551–563.

Hayashi K, Matsui Y. 2006. Meisetz, a novel histone tri-methyltransferase, regulates meiosis-specific epigenesis. Cell Cycle Georget Tex 5: 615–620.

Helleboid P-Y, Heusel M, Duc J, Piot C, Thorball CW, Coluccio A, Pontis J, Imbeault M, Turelli P, Aebersold R, et al. 2019. The interactome of KRAB zinc finger proteins reveals the evolutionary history of their functional diversification. EMBO J 38: e101220.

Huntley S, Baggott DM, Hamilton AT, Tran-Gyamfi M, Yang S, Kim J, Gordon L, Branscomb E, Stubbs L. 2006. A comprehensive catalog of human KRAB-associated zinc finger genes: insights into the evolutionary history of a large family of transcriptional repressors. Genome Res 16: 669–677.

Imbeault M, Helleboid P-Y, Trono D. 2017. KRAB zinc-finger proteins contribute to the evolution of gene regulatory networks. Nature 543: 550–554.

Iouranova A, Grun D, Rossy T, Duc J, Coudray A, Imbeault M, de Tribolet-Hardy J, Turelli P, Persat A, Trono D. 2022. KRAB zinc finger protein ZNF676 controls the transcriptional influence of LTR12-related endogenous retrovirus sequences. Mob DNA 13: 4.

Jacobs FMJ, Greenberg D, Nguyen N, Haeussler M, Ewing AD, Katzman S, Paten B, Salama SR, Haussler D. 2014. An evolutionary arms race between KRAB zinc-finger genes *ZNF91/93* and SVA/L1 retrotransposons. Nature 516: 242–245.

Johansson PA, Brattås PL, Douse CH, Hsieh P, Adami A, Pontis J, Grassi D, Garza R, Sozzi E, Cataldo R, et al. 2022. A cis-acting structural variation at the ZNF558 locus controls a gene regulatory network in human brain development. Cell Stem Cell 29: 52–69.e8.

Karczewski KJ, Francioli LC, Tiao G, Cummings BB, Alföldi J, Wang Q, Collins RL, Laricchia KM, Ganna A, Birnbaum DP, et al. 2020. The mutational constraint spectrum quantified from variation in 141,456 humans. Nature 581: 434–443.

Katoh K, Standley DM. 2013. MAFFT Multiple Sequence Alignment Software Version 7: Improvements in Performance and Usability. Mol Biol Evol 30: 772–780.

Kim S, Cho C-S, Han K, Lee J. 2016. Structural Variation of Alu Element and Human Disease. Genomics Inform 14: 70–77.

Kojima KK. 2018. Human transposable elements in Repbase: genomic footprints from fish to humans. Mob DNA 9: 2.

Langmead B, Salzberg SL. 2012. Fast gapped-read alignment with Bowtie 2. Nat Methods 9: 357–359.

Lawrence M, Carey V, Gentleman R. 2022. rtracklayer: R interface to genome annotation files and the UCSC genome browser. https://bioconductor.org/packages/rtracklayer/.

Lek M, Karczewski KJ, Minikel EV, Samocha KE, Banks E, Fennell T, O’Donnell-Luria AH, Ware JS, Hill AJ, Cummings BB, et al. 2016. Analysis of protein-coding genetic variation in 60,706 humans. Nature 536: 285–291.

Li X, Ito M, Zhou F, Youngson N, Zuo X, Leder P, Ferguson-Smith AC. 2008. A Maternal-Zygotic Effect Gene, Zfp57, Maintains Both Maternal and Paternal Imprints. Dev Cell 15: 547–557.

Liu H, Chang L-H, Sun Y, Lu X, Stubbs L. 2014. Deep vertebrate roots for mammalian zinc finger transcription factor subfamilies. Genome Biol Evol 6: 510–525.

Lukic S, Nicolas J-C, Levine AJ. 2014. The diversity of zinc-finger genes on human chromosome 19 provides an evolutionary mechanism for defense against inherited endogenous retroviruses. Cell Death Differ 21: 381–387.

McLaren PJ, Gawanbacht A, Pyndiah N, Krapp C, Hotter D, Kluge SF, Götz N, Heilmann J, Mack K, Sauter D, et al. 2015. Identification of potential HIV restriction factors by combining evolutionary genomic signatures with functional analyses. Retrovirology 12: 41.

McLaren W, Gil L, Hunt SE, Riat HS, Ritchie GRS, Thormann A, Flicek P, Cunningham F. 2016. The Ensembl Variant Effect Predictor. Genome Biol 17: 122.

Najafabadi HS, Albu M, Hughes TR. 2015. Identification of C2H2-ZF binding preferences from ChIP-seq data using RCADE. Bioinformatics 31: 2879–2881.

Najafabadi HS, Garton M, Weirauch MT, Mnaimneh S, Yang A, Kim PM, Hughes TR. 2017. Non-base-contacting residues enable kaleidoscopic evolution of metazoan C2H2 zinc finger DNA binding. Genome Biol 18: 167.

Nowick K, Gernat T, Almaas E, Stubbs L. 2009. Differences in human and chimpanzee gene expression patterns define an evolving network of transcription factors in brain. Proc Natl Acad Sci 106: 22358–22363.

Ono M, Kawakami M, Takezawa T. 1987. A novel human nonviral retroposon derived from an endogenous retrovirus. Nucleic Acids Res 15: 8725–8737.

Ozata DM, Gainetdinov I, Zoch A, O’Carroll D, Zamore PD. 2019. PIWI-interacting RNAs: small RNAs with big functions. Nat Rev Genet 20: 89–108.

Partridge EC, Chhetri SB, Prokop JW, Ramaker RC, Jansen CS, Goh S-T, Mackiewicz M, Newberry KM, Brandsmeier LA, Meadows SK, et al. 2020. Occupancy maps of 208 chromatin-associated proteins in one human cell type. Nature 583: 720–728.

Patoori S, Barnada SM, Large C, Murray JI, Trizzino M. 2022. Young transposable elements rewired gene regulatory networks in human and chimpanzee hippocampal intermediate progenitors. Development 149: dev200413.

Playfoot CJ, Duc J, Sheppard S, Dind S, Coudray A, Planet E, Trono D. 2021. Transposable elements and their KZFP controllers are drivers of transcriptional innovation in the developing human brain. Genome Res 31: 1531–1545.

Playfoot CJ, Sheppard S, Planet E, Trono D. 2022. Transposable elements contribute to the spatiotemporal microRNA landscape in human brain development. RNA N Y N 28: 1157–1171.

Pontis J, Planet E, Offner S, Turelli P, Duc J, Coudray A, Theunissen TW, Jaenisch R, Trono D. 2019. Hominoid-Specific Transposable Elements and KZFPs Facilitate Human Embryonic Genome Activation and Control Transcription in Naive Human ESCs. Cell Stem Cell 24: 724–735.e5.

Pontis J, Pulver C, Playfoot CJ, Planet E, Grun D, Offner S, Duc J, Manfrin A, Lutolf MP, Trono D. 2022. Primate-specific transposable elements shape transcriptional networks during human development. Nat Commun 13: 7178.

Quenneville S, Verde G, Corsinotti A, Kapopoulou A, Jakobsson J, Offner S, Baglivo I, Pedone PV, Grimaldi G, Riccio A, et al. 2011. In embryonic stem cells, ZFP57/KAP1 recognize a methylated hexanucleotide to affect chromatin and DNA methylation of imprinting control regions. Mol Cell 44: 361–372.

Ramírez F, Ryan DP, Grüning B, Bhardwaj V, Kilpert F, Richter AS, Heyne S, Dündar F, Manke T. 2016. deepTools2: a next generation web server for deep-sequencing data analysis. Nucleic Acids Res 44: W160–W165.

Schmitges FW, Radovani E, Najafabadi HS, Barazandeh M, Campitelli LF, Yin Y, Jolma A, Zhong G, Guo H, Kanagalingam T, et al. 2016. Multiparameter functional diversity of human C2H2 zinc finger proteins. Genome Res 26: 1742–1752.

Seczynska M, Bloor S, Cuesta SM, Lehner PJ. 2021. Genome surveillance by HUSH-mediated silencing of intronless mobile elements. Nature 1–9.

Takahashi N, Coluccio A, Thorball CW, Planet E, Shi H, Offner S, Turelli P, Imbeault M, Ferguson-Smith AC, Trono D. 2019. ZNF445 is a primary regulator of genomic imprinting. Genes Dev 33: 49–54.

Trono D. 2015. Transposable Elements, Polydactyl Proteins, and the Genesis of Human-Specific Transcription Networks. Cold Spring Harb Symp Quant Biol 80: 281–288.

Turelli P, Playfoot C, Grun D, Raclot C, Pontis J, Coudray A, Thorball C, Duc J, Pankevich EV, Deplancke B, et al. 2020. Primate-restricted KRAB zinc finger proteins and target retrotransposons control gene expression in human neurons. Sci Adv 6: eaba3200.

Tycko J, DelRosso N, Hess GT, Aradhana, Banerjee A, Mukund A, Van MV, Ego BK, Yao D, Spees K, et al. 2020. High-Throughput Discovery and Characterization of Human Transcriptional Effectors. Cell 183: 2020–2035.e16.

Urrutia R. 2003. KRAB-containing zinc-finger repressor proteins. Genome Biol 4: 231.

Venkataraman A, Yang K, Irizarry J, Mackiewicz M, Mita P, Kuang Z, Xue L, Ghosh D, Liu S, Ramos P, et al. 2018. A toolbox of immunoprecipitation-grade monoclonal antibodies to human transcription factors. Nat Methods 15: 330–338.

Wagner S, Hess MA, Ormonde-Hanson P, Malandro J, Hu H, Chen M, Kehrer R, Frodsham M, Schumacher C, Beluch M, et al. 2000. A broad role for the zinc finger protein ZNF202 in human lipid metabolism. J Biol Chem 275: 15685–15690.

Wang H, Xing J, Grover D, Hedges DJ, Han K, Walker JA, Batzer MA. 2005. SVA Elements: A Hominid-specific Retroposon Family. J Mol Biol 354: 994–1007.

Wang W, Shang W, Zou J, Liu K, Liu M, Qiu X, Zhang H, Wang K, Wang N. 2022. ZNF667 facilitates angiogenesis after myocardial ischemia through transcriptional regulation of VASH1 and Wnt signaling pathway. Int J Mol Med 50: 129.

Weirauch MT, Yang A, Albu M, Cote AG, Montenegro-Montero A, Drewe P, Najafabadi HS, Lambert SA, Mann I, Cook K, et al. 2014. Determination and inference of eukaryotic transcription factor sequence specificity. Cell 158: 1431–1443.

Wolf D, Goff SP. 2009. Embryonic stem cells use ZFP809 to silence retroviral DNAs. Nature 458: 1201–1204.

Xiang X, Tao Y, DiRusso J, Hsu F-M, Zhang J, Xue Z, Pontis J, Trono D, Liu W, Clark AT. 2022. Human reproduction is regulated by retrotransposons derived from ancient Hominidae-specific viral infections. Nat Commun 13: 463.

Yan J, Enge M, Whitington T, Dave K, Liu J, Sur I, Schmierer B, Jolma A, Kivioja T, Taipale M, et al. 2013. Transcription factor binding in human cells occurs in dense clusters formed around cohesin anchor sites. Cell 154: 801–813.

Yang P, Wang Y, Hoang D, Tinkham M, Patel A, Sun M-A, Wolf G, Baker M, Chien H-C, Lai K-YN, et al. 2017a. A placental growth factor is silenced in mouse embryos by the zinc finger protein ZFP568. Science 356: 757–759.

Yang P, Wang Y, Macfarlan TS. 2017b. The Role of KRAB-ZFPs in Transposable Element Repression and Mammalian Evolution. Trends Genet 33: 871–881.

Zeng Y, Wang W, Ma J, Wang X, Guo M, Li W. 2012. Knockdown of ZNF268, which Is Transcriptionally Downregulated by GATA-1, Promotes Proliferation of K562 Cells. PLOS ONE 7: e29518.

Zhang Y, Liu T, Meyer CA, Eeckhoute J, Johnson DS, Bernstein BE, Nusbaum C, Myers RM, Brown M, Li W, et al. 2008. Model-based Analysis of ChIP-Seq (MACS). Genome Biol 9: R137.

