## Supplemental_Figures for "Genetic features and genomic targets of human KRAB-Zinc Finger Proteins"

### Supplementary Figures

**Figure S1**

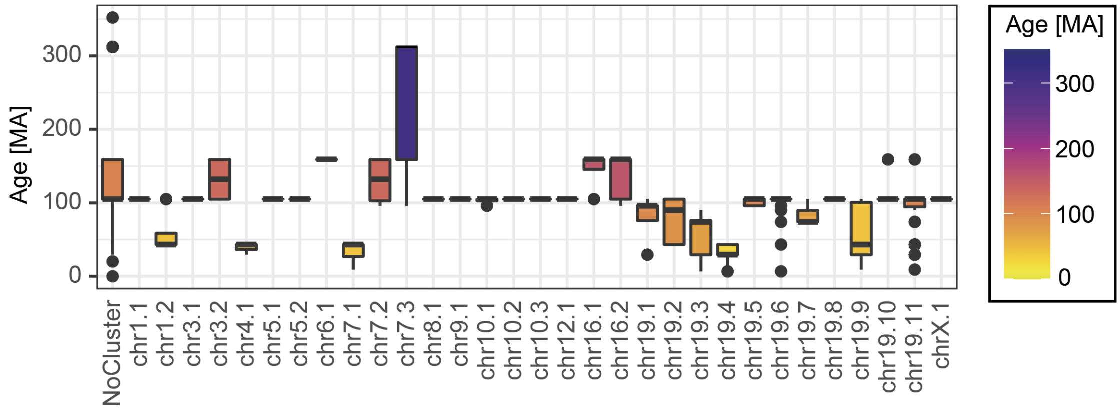


#### Figure S1: Age and polymorphism of KZFPs in clusters

A) Boxplots of the ages per cluster as defined in Figure 1, coloured by the median age of the KZFPs in the clusters.

**Figure S2**

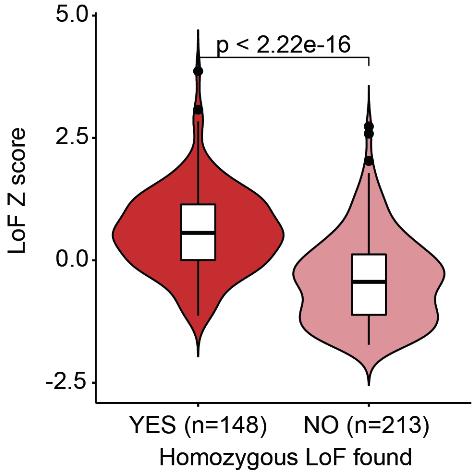


#### Figure S2: Relationship between LoF constraint and presence of homozygous LoF variants.

KZFPs in which homozygous LoF variants were found are less constrained than KZFPs were only heterozygous LoF variants were identified. The p-value were obtained with the Wilcoxon rank sum test.

#### Figure S3

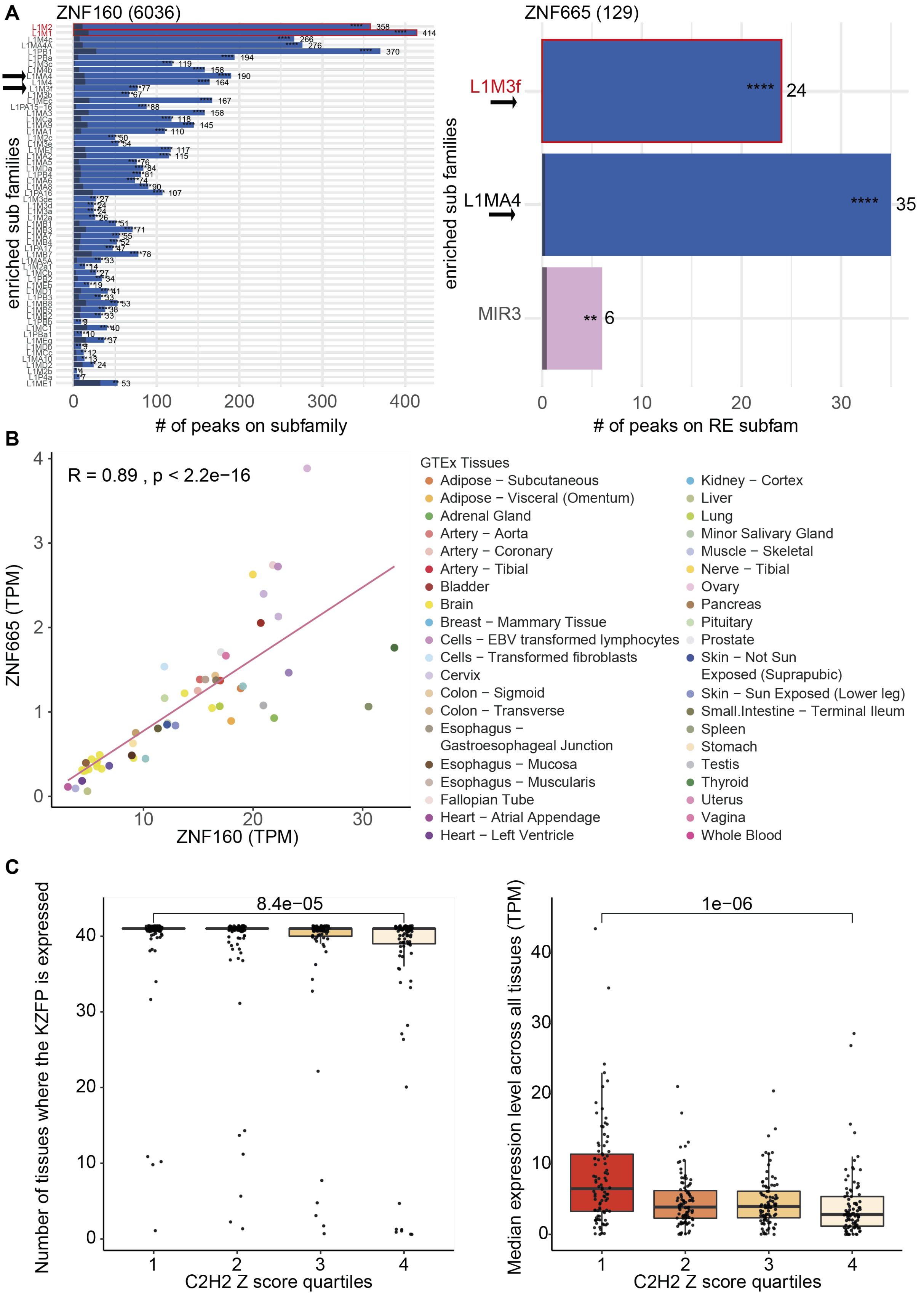
A
ZNF160 (6036)
enriched sub families
### of peaks on subfamily
ZNF665 (129)
L1M3f
L1MA4
MIR3
enriched sub families
### of peaks on RE subfam
B
R = 0.89 , p < 2.2e−16
ZNF665 (TPM)
ZNF160 (TPM)
GTEx Tissues
Adipose − Subcutaneous
Adipose − Visceral (Omentum)
Adrenal Gland
Artery − Aorta
Artery − Coronary
Artery − Tibial
Bladder
Brain
Breast − Mammary Tissue
Cells − EBV transformed lymphocytes
Cells − Transformed fibroblasts
Cervix
Colon − Sigmoid
Colon − Transverse
Esophagus − Gastroesophageal Junction
Esophagus − Mucosa
Esophagus − Muscularis
Fallopian Tube
Heart − Atrial Appendage
Heart − Left Ventricle
Kidney − Cortex
Liver
Lung
Minor Salivary Gland
Muscle − Skeletal
Nerve − Tibial
Ovary
Pancreas
Pituitary
Prostate
Skin − Not Sun Exposed (Suprapubic)
Skin − Sun Exposed (Lower leg)
Small.Intestine − Terminal Ileum
Spleen
Stomach
Testis
Thyroid
Uterus
Vagina
Whole Blood
C
8.4e−05
Number of tissues where the KZFP is expressed
C2H2 Z score quartiles
1e−06
Median expression level across all tissues (TPM)
C2H2 Z score quartiles

#### Figure S3: Comparison of ZNF160 and ZNF665 paraloges.

A) Similar genomic targets for ZNF160 and ZNF665. Enrichments of ZNF160 and ZNF665 peaks over different repetitive element subfamilies (FDR < 0.01). The width of the coloured bars represents the number of peaks per subfamily also shown as a number on the right of the bar. The black transparent bars represent the expected number of peaks following a random distribution. The FDR of the enrichment is shown with stars (FDR < 0.0001 = ****, < 0.001=***, < 0.01=**, < 0.05=*, >= 0.05 = n.s). The y-axis is ordered by FDR. The number next to the title indicates the total number of peaks for the experiment. Primary targets for each KZFP are highlighted in red. (B) Expression levels in transcripts per million (TPM) of ZNF160 and ZNF665 across all tissues depicted in GTEx (https://gtexportal.org). C) Biological consequences of KZFP constraint. Left: KZFPs in the highest quartile of C2H2 constraint are expressed in more tissues than KZFPs in the lowest quartile. Right: The mean expression levels of KZFPs in the first quartile of C2H2 constraint is higher than for those KZFPs in the fourth constraint quartile (lowest constraint). All p-values are from Wilcoxon rank sum tests.

**Figure S4**

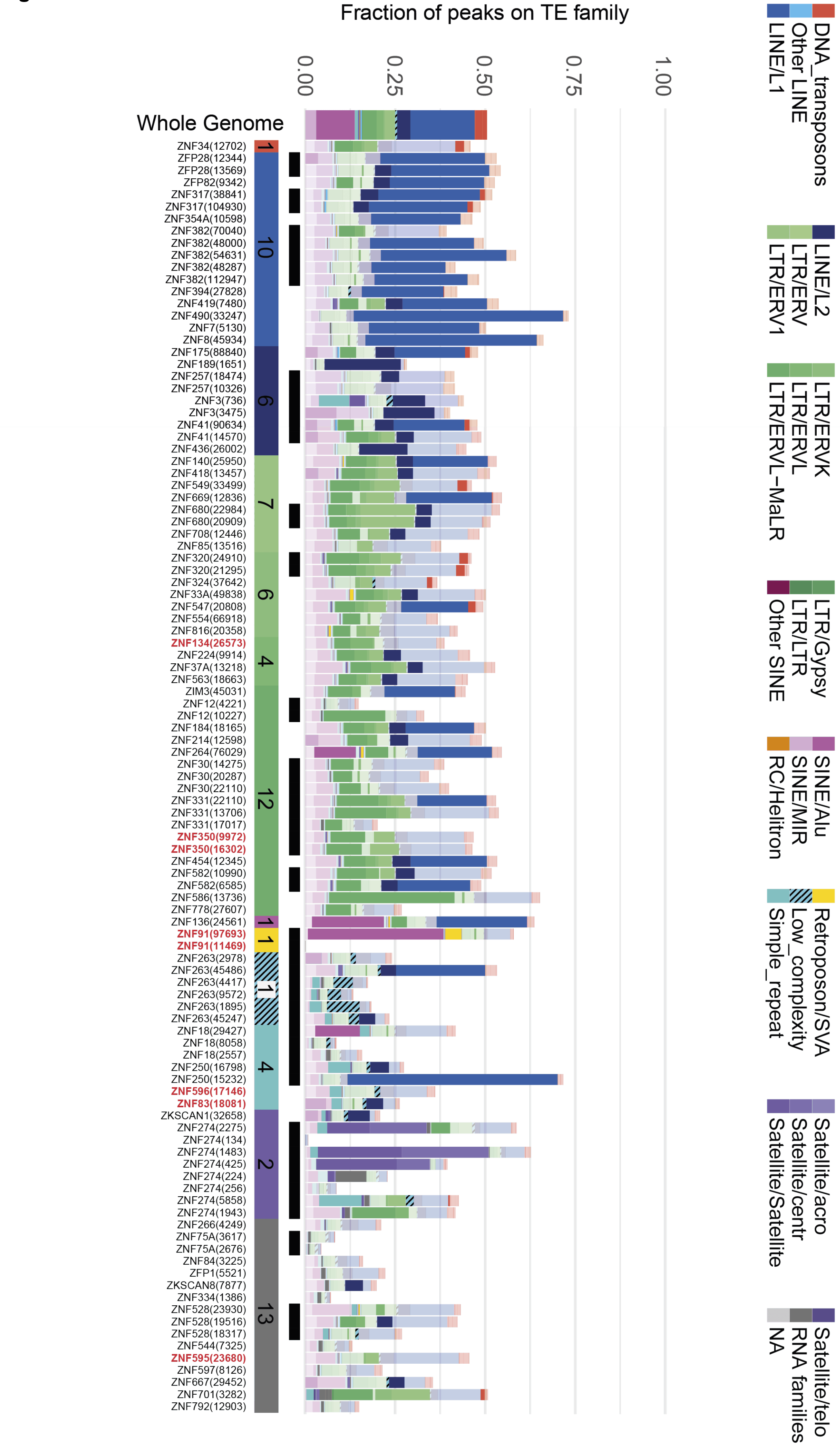
Fraction of peaks on TE family
0.00
0.25
0.50
0.75
1.00
Whole Genome
ZNF34(12702)
ZFP28(12344)
ZFP28(13569)
ZFP82(9342)
ZNF317(38841)
ZNF317(104930)
ZNF354A(10598)
ZNF382(70040)
ZNF382(48000)
ZNF382(54631)
ZNF382(48287)
ZNF382(112947)
ZNF394(27828)
ZNF419(7480)
ZNF490(33247)
ZNF7(5130)
ZNF8(45934)
ZNF175(88840)
ZNF189(1651)
ZNF257(18474)
ZNF257(10326)
ZNF3(736)
ZNF3(3475)
ZNF41(90634)
ZNF41(14570)
ZNF436(26002)
ZNF140(25950)
ZNF418(13457)
ZNF549(33499)
ZNF669(12836)
ZNF680(22984)
ZNF680(20909)
ZNF708(12446)
ZNF85(13516)
ZNF320(24910)
ZNF320(21295)
ZNF324(37642)
ZNF33A(49838)
ZNF547(20808)
ZNF554(66918)
ZNF816(20358)
ZNF134(26573)
ZNF224(9914)
ZNF37A(13218)
ZNF563(18663)
ZIM3(45031)
ZNF12(4221)
ZNF12(10227)
ZNF184(18165)
ZNF214(12598)
ZNF264(76029)
ZNF30(14275)
ZNF30(20287)
ZNF30(22110)
ZNF331(22110)
ZNF331(13706)
ZNF331(17017)
ZNF350(9972)
ZNF350(16302)
ZNF454(12345)
ZNF582(10990)
ZNF582(6585)
ZNF586(13736)
ZNF778(27607)
ZNF136(24561)
ZNF91(97693)
ZNF91(11469)
ZNF263(2978)
ZNF263(45486)
ZNF263(4417)
ZNF263(9572)
ZNF263(1895)
ZNF263(45247)
ZNF18(29427)
ZNF18(8058)
ZNF18(2557)
ZNF250(16798)
ZNF250(15232)
ZNF596(17146)
ZNF83(18081)
ZKSCAN1(32658)
ZNF274(2275)
ZNF274(134)
ZNF274(1483)
ZNF274(425)
ZNF274(224)
ZNF274(256)
ZNF274(5858)
ZNF274(1943)
ZNF266(4249)
ZNF75A(3617)
ZNF75A(2676)
ZNF84(3225)
ZFP1(5521)
ZKSCAN8(7877)
ZNF334(1386)
ZNF528(23930)
ZNF528(19516)
ZNF528(18317)
ZNF544(7325)
ZNF595(23680)
ZNF597(8126)
ZNF667(29452)
ZNF701(3282)
ZNF792(12903)
1
10
9
7
6
4
12
1
1
1
4
2
13
DNA_transposons
Other LINE
LINE/L1
LINE/L2
LTR/ERV
LTR/ERV1
LTR/ERVK
LTR/ERVL
LTR/ERVL-MaLR
LTR/Gypsy
LTR/LTR
Other SINE
SINE/Alu
SINE/MIR
RC/Helitron
Retroposon/SVA
Low_complexity
Simple_repeat
Satellite/acro
Satellite/centr
Satellite/Satellite
Satellite/telo
RNA families
NA

#### Figure S4: Peaks, targets identified with external data and ages of KZFPs relative to the ages of their targets

A) Bar graph showing the fraction of peaks over repetitive element (RE) families for experiments conducted using different over-expression protocols (Table S2). Columns are ordered by the most enriched family which are indicated by the horizontal bar below, along with the number of KZFPs for each category. Replicate experiments are indicated by black squares above the horizontal bar. Significant enrichments (FDR < 0.05) are show in fully opaque colours where non-significant enrichments are transparent. The leftmost bar shows the genome occupancy of all RE families. The total number of peaks per experiment is indicated in brackets after the KZFP name below each bar. KZFPs that are not also represented in Figure 5A are highlighted in red.

**Figure S5**

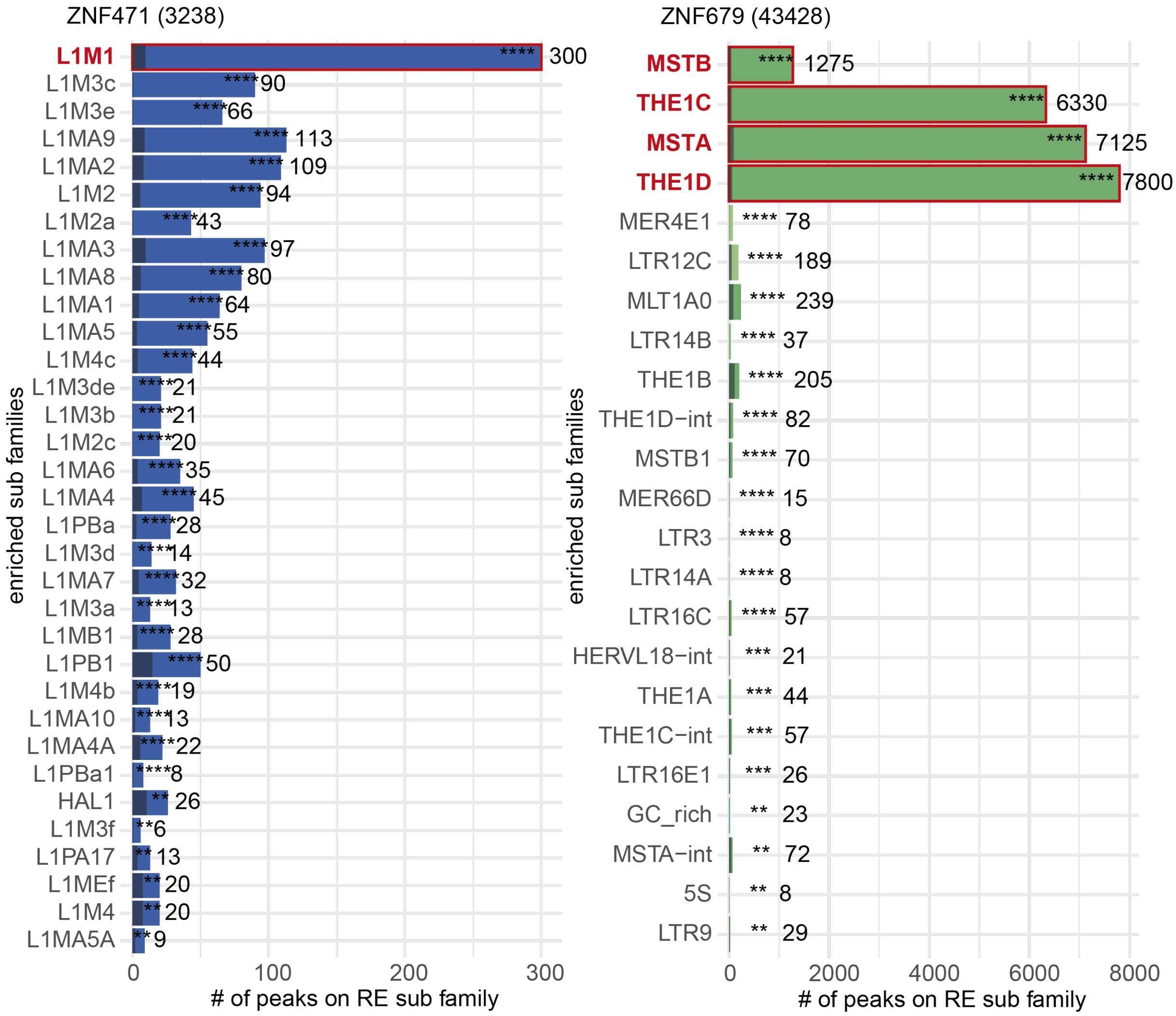


#### Figure S5: Example of Primary vs. Secondary targets

Enrichments of ZNF471 and ZNF679 peaks over different repetitive element subfamilies (FDR < 0.01). Primary targets for each KZFP are highlighted in red. The width of the coloured bars represents the number of peaks per subfamily also shown as a number on the right of the bar. The black transparent bars represent the expected number of peaks following a random distribution. The FDR of the enrichment is shown with stars (FDR < 0.0001 = ****, < 0.001=***, < 0.01=**, < 0.05=*, >= 0.05 = n.s). The y-axis is ordered by FDR. The number next to the title indicates the total number of peaks for the experiment.

**Figure S6**

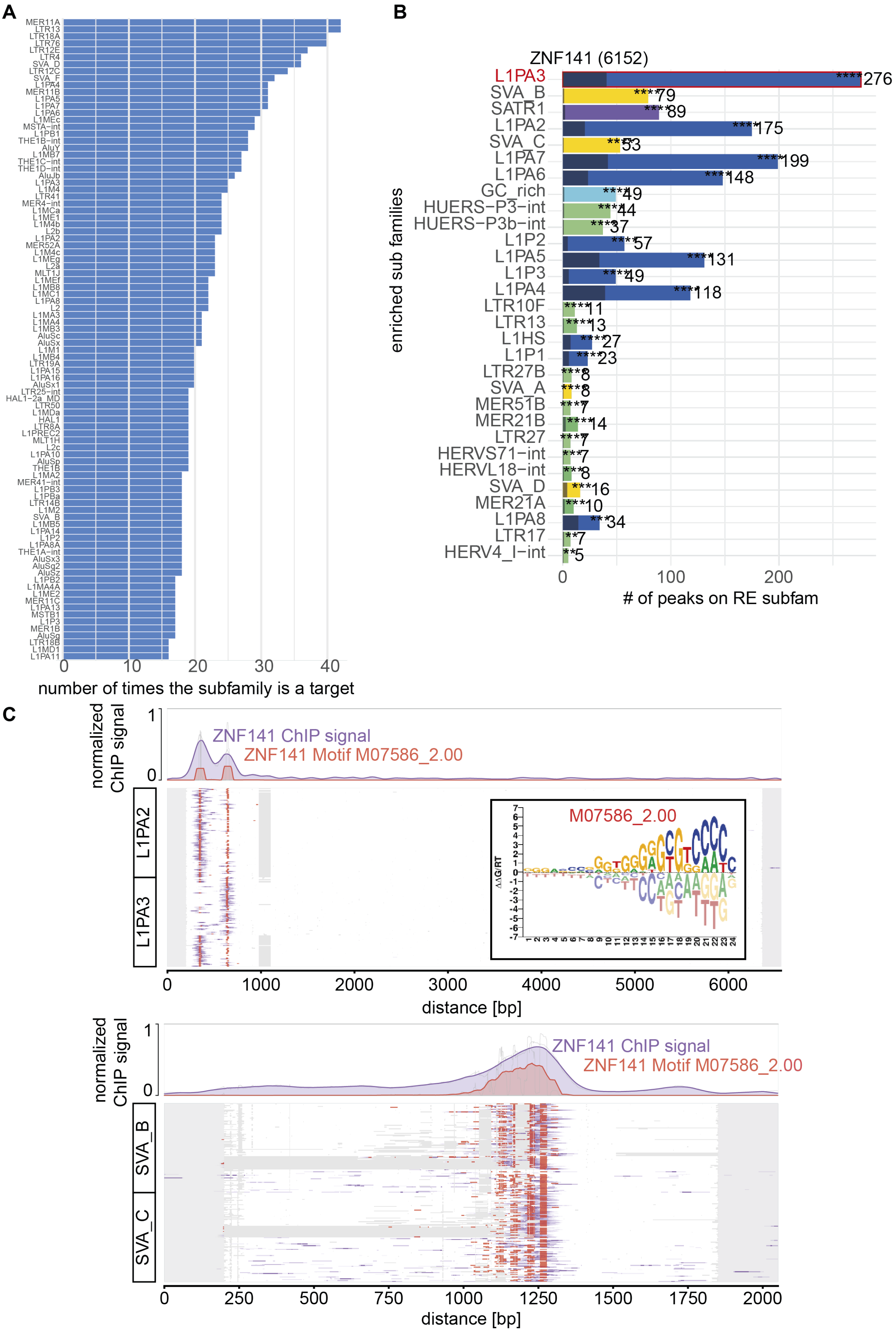
A
B
C
number of times the subfamily is a target
0
10
20
30
40
ZNF141 (6152)
enriched sub families
L1PA3
SVA_B
SATR1
L1PA2
SVA_C
L1PA7
L1PA6
GC_rich
HUERS−P3−int
HUERS−P3b−int
L1P2
L1PA5
L1P3
L1PA4
LTR10F
LTR13
L1HS
L1P1
LTR27B
SVA_A
MER51B
MER21B
LTR27
HERVS71−int
HERVL18−int
SVA_D
MER21A
L1PA8
LTR17
HERV4_I−int
****276
****79
****89
****175
****53
****199
****148
****49
****44
****37
****57
****131
****49
****118
****11
****13
****27
****23
***8
***8
****7
****14
****7
***7
***8
***16
***10
***34
**7
**5
0
100
200
### of peaks on RE subfam
normalized ChIP signal
1
0
ZNF141 ChIP signal
ZNF141 Motif M07586_2.00
L1PA2
L1PA3
M07586_2.00
0
1000
2000
3000
4000
5000
6000
distance [bp]
normalized ChIP signal
1
0
ZNF141 ChIP signal
ZNF141 Motif M07586_2.00
SVA_B
SVA_C
0
250
500
750
1000
1250
1500
1750
2000
distance [bp]

#### Figure S6: ZNF141 is binding to SVA VNTR

A) Bar graph with the number of KZFPs enriched on TE subfamilies (rows). TE subfamilies with more than 15 enriched KZFPs with an FDR < 0.05 are shown. B) Enrichment of ZNF141 peaks over different repetitive element subfamilies (FDR < 0.01). The width of the coloured bars represents the number of peaks per subfamily also shown as a number on the right of the bar. The black transparent bars represent the expected number of peaks following a random distribution. The FDR of the enrichment is shown with stars (FDR < 0.0001 = ****, < 0.001=***, < 0.01=**, < 0.05=*, >= 0.05 = n.s). The y-axis is ordered by FDR. The number next to the title indicates the total number of peaks for the experiment.C) Multiple sequence alignment (MSA) over the most enriched targets L1PA2 and 3 (top) and SVA_B and C in (bottom). Up to 200 elements for the indicated targets where aligned, selecting first elements overlapping with a peak and then the longest elements. The signal of the ZNF141 ChIP was laid over the alignment in purple. The locations of the motif identified in (Weirauch et al. 2014) is shown in red. The motif is shown on the right of the top panel- The average signal normalized for each element (row-wise) can be seen as a line plot above the MSA plots.

**Figure S7**

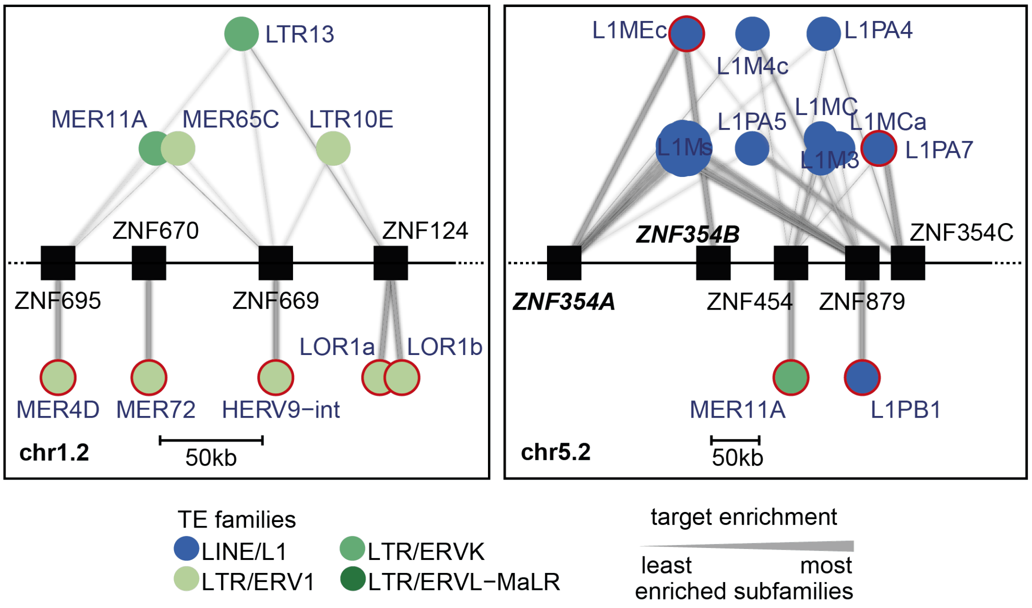

##### Figure S7: More examples of shared targets

A) Networks for clusters chr1.2 and chr5.2 were targets (circles) of each KZFP (squares) are shown as connected edges and the amount of binding is represented by the line thickness. The thickest line for each KZFP represents the TE subfamily with the highest -log10(FDR) and then scales linearly to the lowest value. For visibility, only the best targets (below) and shared targets (above) are shown. The TE subfamilies are coloured according to their families. Primary targets for each KZFP are highlighted in red.

#### Tables

##### Table S1: Census of human KZFPs

Locations and annotations of all human KRAB and Zinc-Finger domain pairs as described in the Methods.

##### Table S2: ChIP-seq data on human KZFPs

Information on the available ChIP-seq data for all human KZFPs.

##### Table S3

Table of primary targets of KZFPs

##### Table S4

Data underlying Figure 5C
